# A pan-cohort transcriptional landscape of breast cancer maps subtype and microenvironmental programs

**DOI:** 10.64898/2026.08.31.748296

**Authors:** Sonali Arora, Ramya Suresh, Nikolas Holland, Gregory Glatzer, Matt Jensen, Eric Q. Konnick, Colin C. Pritchard, Yi Li, Heather A. Parsons, Sara A. Hurvitz, Eric C Holland

## Abstract

Breast cancer comprises heterogeneous transcriptional states that are incompletely captured by discrete clinical or molecular subtype labels. To visualize this heterogeneity in a unified framework, we integrated bulk RNA-seq data from 2,284 patient samples across 13 studies using 18,089 protein coding genes, a harmonized processing pipeline, batch correction, consensus clustering and PaCMAP dimensionality reduction to construct an interactive breast cancer transcriptional landscape. Consensus clustering identified five major regions, which were annotated using PAM50 scores calculated for each sample: Luminal A, Luminal B, HER2-enriched, and two basal-associated clusters. The basal clusters separated into an immune-rich region marked by T cell–inflamed, tumor-associated macrophages (TAM), and low-purity signatures, and a cell-cycle–driven region enriched for proliferation and DNA replication programs. Overlay of marker genes, pathways, kinases, neuronal-like signaling programs, cancer associated fibroblasts (CAF) states, and TAM programs revealed spatially organized subtype biology and microenvironmental heterogeneity. Finally, projection of therapy-associated resistance signatures identified landscape regions linked to predicted resistance to HER2-targeted therapy and hormone receptor–directed endocrine therapies. By enabling interactive exploration of transcriptional states, marker genes, pathways, and therapeutic response programs, this resource provides a community framework for biomarker discovery in breast cancer.

**Significance statement:** We present a unified, multi-cohort transcriptional landscape of breast cancer that organizes canonical subtypes along continuous biological axes and reveals spatially structured regions of therapeutic sensitivity and resistance. By enabling projection of patient samples and patient-derived models into this framework, we provide a practical tool for interpreting tumor biology and guiding translational discovery.

**Graphical abstract caption:** We integrate 13 breast cancer transcriptomic datasets using a harmonized processing pipeline with batch correction and dimensionality reduction to construct a unified landscape spanning basal, HER2-enriched, luminal A, luminal B, and normal-like subtypes. Continuous gradients of proliferation, endocrine signaling, metabolic activity, and tissue architecture organize the space and recapitulate subtype biology. Overlay of clinical outcomes reveals high-risk regions beyond discrete subtype boundaries, while integration of gene expression and copy number alterations links genomic events to pathway-level programs. Microenvironmental features, including cancer-associated fibroblast programs, map to distinct regions of the landscape. Projection of drug response signatures identifies zones of sensitivity and intrinsic resistance. The framework supports embedding of patient samples and patient-derived xenograft models and is available as an interactive resource for exploration and projection of new datasets.

**One Sentence Summary:** A landscape built using only transcriptomic analysis for breast cancer reveals novel insights about subtype-specific biology.

## INTRODUCTION

Breast cancer is a heterogeneous disease comprising multiple clinically and molecularly distinct subtypes that differ in prognosis, therapeutic response, and underlying biology^1^. Despite substantial advances in genomic and transcriptomic profiling, patient stratification in clinical practice remains largely anchored to a limited set of biomarkers, including hormone receptor status and ERBB2 amplification. While these classifications have enabled targeted therapies, they incompletely capture the full spectrum of transcriptional diversity observed across tumors. As a result, patients with similar clinical annotations can exhibit markedly different outcomes^2,3^, underscoring the need for frameworks that more comprehensively resolve tumor heterogeneity.

Transcriptional subtyping has defined intrinsic breast cancer classes, including basal-like, HER2-enriched, luminal A, luminal B, and normal-like tumors^4–7^. These subtypes reflect major biological axes, including proliferation, endocrine signaling, and growth factor activation. However, increasing evidence^8^ suggests that these categories represent coarse partitions of a more continuous transcriptional landscape. Tumors often exhibit features that span subtype boundaries, and intermediate states may carry clinically relevant information that is not captured by discrete labels.

In prior studies of brain^9–11^ and lung^12^ cancers, dimensionality-reduced representations of large publicly-available patient tumor transcriptomic datasets have been constructed and used to preserve global structure while enabling comparison across samples. These transcriptional landscapes, built using all the protein coding genes in the human genome, can reveal coherent biological programs, identify previously unrecognized clusters, and map clinically relevant features onto continuous spaces. These approaches provide a unifying framework for integrating heterogeneous datasets and for interpreting tumor states beyond predefined subtype classifications.

Here, we extend this framework to breast cancer by integrating breast tumors from 13 independent transcriptomic datasets, processed through a harmonized bioinformatic pipeline. Following batch correction, clustering and dimensionality reduction, we construct a unified transcriptional landscape spanning basal, HER2-enriched, luminal A, luminal B, and normal-like tumors. We project clinical annotations, genomic alterations, and pathway-level programs onto this space to resolve subtype structure and uncover continuous biological axes. In addition, we map microenvironmental features, including cancer-associated fibroblast programs and innervation, and evaluate how drug response signatures distribute across the landscape.

While single-cell atlases^8^ have provided important insights into tumor heterogeneity, they remain limited by cohort size and cost. By integrating and reanalyzing publicly available bulk RNA-seq data from 2,284 patient samples, we leverage the vast sample size to resolve continuous transcriptional structure across breast cancer cases. The resulting Oncoscape(<u>Breast Landscapes</u>) resource enables interactive visualization, projection of new datasets, and interrogation of genes and pathways across a unified breast cancer landscape.

## RESULTS

### Construction of a breast cancer transcriptomic landscape

We integrated raw sequencing data from 2,284 patient samples across 13 independent studies and processed all datasets using a harmonized bioinformatic pipeline. The integrated cohort included 2,284 breast cancer RNA-seq samples from 13 publicly available datasets: AURORA^13^, CMI-MBC^14^, TCGA-BRCA^15^, Fudan University Shanghai Cancer Center (FUSCC) triple-negative breast cancer cohort^16^, and other smaller clinically well annotated studies from GEO^17–24^ (Table S1).

Following quantification of 18,089 protein-coding gene expression, raw gene counts were batch-corrected and variance-stabilized to enable cross-cohort integration, which were used for three complementary analyses. First, consensus clustering was performed using 18,089 protein-coding genes to identify stable transcriptional groups, yielding five major clusters across the integrated cohort (Figure S1a). A smaller group of 56 samples localized away from the main embedding and, although not robustly recovered as an independent consensus cluster, showed distinct downstream transcriptional and genomic features; we therefore retained it as cluster F for separate interpretation. Second, we computed Gene set variation analysis (GSVA) scores for PAM50 gene sets^4^ for all samples (Table S2), enabling subtype annotation across datasets regardless of whether PAM50 labels were reported in the original metadata. Third, we compared multiple dimensionality-reduction methods, including Principal Component Analysis (PCA), t-distributed Stochastic Neighbor Embedding (t-SNE), uniform manifold approximation and projection (UMAP), and Pairwise controlled manifold approximation (PaCMAP)^25^ to visualize the structure of the integrated cohort (Figure S1b). Clustering was used for cluster discovery, whereas dimensionality-reduction methods were used only for visualization. Because PaCMAP provided the clearest separation of dataset-integrated, PAM50-aligned, and consensus cluster-defined regions, it was selected as the final visualization framework for the breast cancer landscape (Figure 1a; Figure S1b).

**Figure 1.**
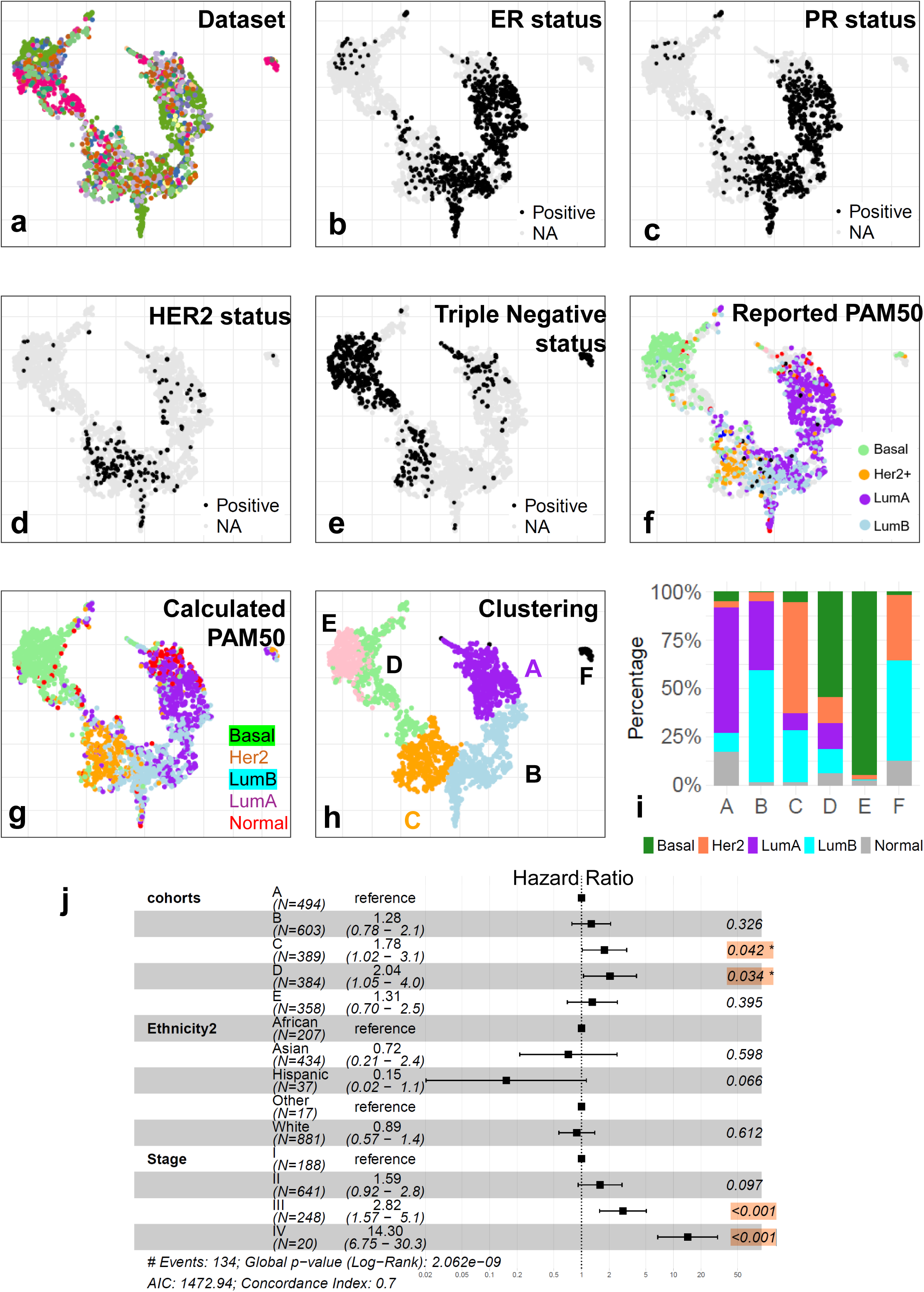
Construction of a unified breast cancer transcriptional landscape. (A) PaCMAP embedding of 2,284 breast cancer samples colored by dataset of origin, showing effective integration and absence of batch-driven clustering. (B-E) Embedding colored by clinical receptor status: estrogen receptor (ER), progesterone receptor (PR), HER2 status, and triple-negative breast cancer (TNBC), showing spatial localization of major clinical phenotypes. (F) Samples colored by reported PAM50 subtype (basal-like, HER2-enriched, luminal A, luminal B, normal-like), revealing distinct subtype-enriched regions. (G) GSVA scores for PAM50 gene signatures projected onto the landscape, showing continuous gradients aligned with subtype regions. (H) consensus clustering using k-means reveals 6 distinct clusters. (I) Barplot showing composition of breast cancer subtypes, based on calculated PAM50 scores, for each cluster. (J) Multivariable Cox regression analysis of evaluating the association between cluster and survival, adjusted for ethnicity and stage at diagnosis.

To further assess the stability of the landscape, we repeated the PaCMAP analysis after random subsampling of the integrated cohort at 25%, 50%, 75%, and 90% of samples. Across all subsampled datasets, the major structure of the embedding was preserved, with similar dataset mixing and consistent localization of calculated PAM50 subtypes and consensus cluster-defined regions (Figure S1c). These analyses indicate that the observed landscape organization is not driven by a small subset of samples and support the robustness of the underlying transcriptional structure.

These clusters showed strong, but not fully exclusive, correspondence with canonical breast cancer subtypes. PAM50 subtype classification assigned cluster A predominantly as Luminal A tumors (67.48%), with a smaller contribution from Luminal B tumors (9.91%), whereas Cluster B was classified predominantly as Luminal B tumors (58.04%), while also containing a substantial fraction of Luminal A tumors (35.98%). Cluster C was enriched for HER2-positive tumors (57.32%) but also contained a large proportion of Luminal B samples (26.9%). Cluster D and E were predominantly basal-like, comprising 54.68% and 95.25% basal tumors, respectively (Figure 1i; Table S3). By contrast, Cluster F, despite being annotated as triple-negative, was classified by PAM50 as a mixture of Luminal B and HER2-positive tumors, further highlighting the transcriptional overlap between clinically defined categories.

Although most tumors in the landscape were annotated as primary breast tumors in the original publications, we included 28 metastatic samples spanning multiple metastatic sites. These samples were distributed across the landscape, but were most concentrated in Cluster C, which contained 13 metastatic samples, including four of the five liver metastases. By comparison, Clusters A and B each contained five metastatic samples, Cluster D contained one, and Cluster E contained four. This distribution suggests that Cluster C may capture transcriptional features enriched among metastatic samples in this cohort, particularly liver metastases, although larger metastatic datasets will be needed to validate site-specific patterns (Figure S1j). Projection of clinical annotations onto the embedding revealed coherent spatial organization of major breast cancer phenotypes. ER-positive, PR-positive, HER2-positive, and triple-negative tumors localized to distinct regions of the landscape (Figure 1b–e, Figure S1d-g). While the majority of samples clustered within their expected domains, a subset localized outside canonical regions, suggesting the presence of intermediate states or potential misclassification based on standard clinical annotations. Overlay of reported and calculated PAM50 subtype labels showed clear localization of basal-like, HER2-enriched, Luminal A, Luminal B, and normal-like tumors to distinct regions of the landscape (Figure 1f-g, Figure S1h-i). Among samples with both reported and calculated PAM50 subtype assignments, concordance was high. Additional clinical variables, including stage, lymph node involvement, age at diagnosis, margin status, and pathological staging (M, N, and T), did not form discrete clusters within the landscape (Figure S1k-t), indicating that these clinical features do not correlate with overall gene expression patterns.

Multivariable Cox regression analysis, stratified by ethnicity and stage at diagnosis, demonstrated that Cluster C and Cluster D were independently associated with poorer survival (Cluster C: HR = 1.78, p = 0.042; Cluster D: HR = 2.04, p = 0.034), with advanced stage III–IV remaining the strongest predictor of adverse outcomes (p < 0.001) (Figure 1j).

### Canonical marker genes validate cluster identity across the landscape

To validate the biological fidelity of the landscape, we overlaid canonical breast cancer marker genes onto the cluster-defined embedding. Basal-associated markers, including KRT5^26^, FOXC1^27^, and SOX10^27^, were strongly enriched in Clusters D and E, the two basal-associated regions of the landscape (Figure S2a), and also in a subset of tumors in cluster A. Cluster C, which was enriched for HER2-positive tumors, showed high expression of ERBB2^28^, GRB7^29^, and PGAP3^29^ (Figure 2a, Figure S2a), consistent with HER2-associated signaling. Cluster A, corresponding predominantly to Luminal A tumors, was characterized by elevated expression of endocrine markers ESR1^30^, PGR (Figure 2b-c), and FOXA1^31^, whereas proliferation-associated genes MKI67^32^, CCNB1^32^, and AURKA^32^ were enriched in cluster B, which corresponds predominantly to Luminal B tumors, as well as cluster D and E. (Figure S2a). Notably, although these markers localized to their expected cluster-enriched regions, their expression formed continuous gradients rather than sharp boundaries, further supporting a model in which subtype-associated programs vary across a continuum of transcriptional states.

**Figure 2.**
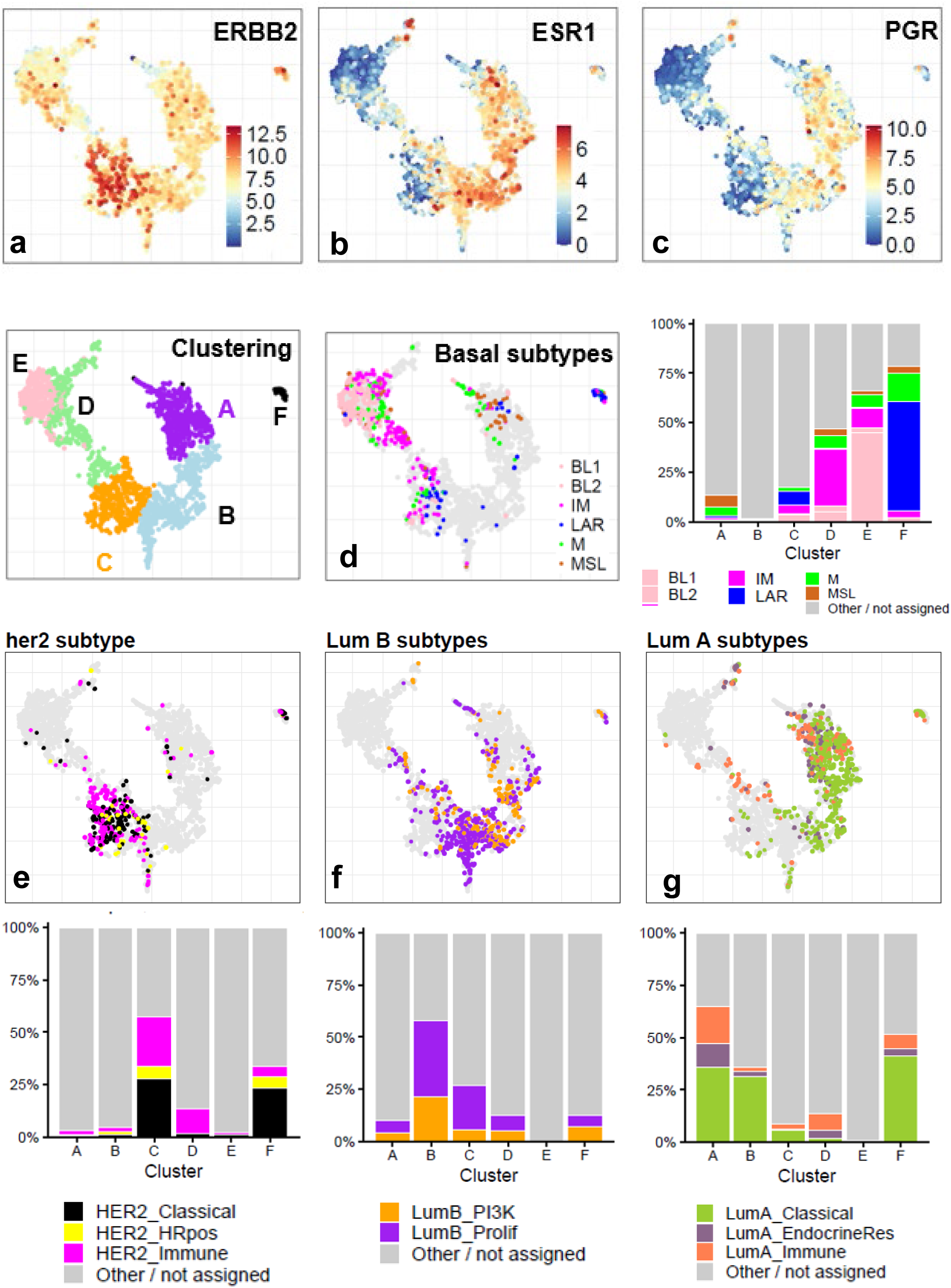
Marker genes and refined subtype programs define regional structure across the breast cancer landscape. (A–C) PaCMAP embeddings colored by expression of key breast cancer marker genes: **ERBB2** (A), **ESR1** (B), and **PGR** (C), showing localization of HER2-associated and hormone receptor–associated transcriptional programs across the landscape. (D) PaCMAP embedding colored by calculated TNBC subtype assignments, accompanied by a stacked bar plot showing the distribution of TNBC subtypes across landscape clusters. (E) PaCMAP embedding colored by calculated HER2 subtype assignments, accompanied by a stacked bar plot showing the distribution of HER2 subtype programs across landscape clusters. (F) PaCMAP embedding colored by calculated Luminal B subtype assignments, accompanied by a stacked bar plot showing the distribution of Luminal B subtype programs across landscape clusters. (G) PaCMAP embedding colored by calculated Luminal A subtype assignments, accompanied by a stacked bar plot showing the distribution of Luminal A subtype programs across landscape clusters.

### Subtype-program concordance reveals structured heterogeneity within canonical breast cancer classes

To further resolve transcriptional heterogeneity within canonical breast cancer classes, we compared landscape clusters with reported and calculated subtype assignments using cluster-by-subtype composition analyses. Among samples with reported Lehmann TNBC subtype^26^ annotations (Figure S2b), subtype distribution was significantly associated with landscape cluster identity (Monte Carlo chi-square p < 1 × 10^-4; Cramér’s V = 0.41; Table S4). Reported LAR tumors localized primarily to Clusters A, B, C, and F, whereas BL1/BL2 tumors were enriched within the basal-associated regions, particularly Cluster E. Consistent with this pattern, calculated TNBC subtype scores (Figure 2d, Figure S2b) across 603 triple-negative tumors showed strong concordance with cluster structure (Monte Carlo chi-square p < 1 × 10^-4; Cramér’s V = 0.45). Cluster D was dominated by the immune-modulatory program (110/181, 60.77%), whereas Cluster E was dominated by BL1 tumors (161/236, 68.22%). Cluster F was enriched for the LAR program (31/44, 70.45%), supporting its interpretation as an AR-enriched, luminal-like TNBC region rather than a canonical basal-like state. Cluster A contained a substantial MSL component among TNBC tumors (29/65, 44.62%), indicating that mesenchymal and stem-like TNBC programs extend into luminal-adjacent regions of the landscape.

We also computed GSVA scores for refined HER2 and luminal subtype programs^28,33,34^(Figure 2e-g, Figure S2b). Calculated HER2 subtype assignments were significantly associated with landscape clusters (Monte Carlo chi-square p < 1 × 10^-4; Cramér’s V = 0.29). Cluster C contained most HER2-classical tumors and was enriched for HER2-classical and HER2-immune programs, whereas Cluster D showed predominant HER2-immune assignment among HER2-subtyped samples (45/52, 86.54%). Luminal subtype programs also showed structured but more diffuse organization. Luminal B subtype composition was associated with cluster identity (Monte Carlo chi-square p = 0.0053; Cramér’s V = 0.16), with Luminal B proliferative programs enriched across Clusters B and C. Luminal A subtype composition showed stronger cluster association (Monte Carlo chi-square p < 1 × 10^-4; Cramér’s V = 0.31), with Luminal A classical tumors enriched in Clusters A and B, while Luminal A immune programs were most prominent among Luminal A-assigned samples in Cluster D (30/52, 57.69%). Together, these analyses show that landscape clusters are concordant with established subtype programs but also reveal cross-subtype transcriptional overlap, including immune-enriched, proliferative, LAR, and mesenchymal programs that distribute across canonical subtype boundaries. TCGA subtype annotations showed a similar spatial organization (Figure S2c), further supporting the biological validity of the landscape.

### Copy-number profiles validate cluster-specific genomic structure

To further validate the cluster-defined landscape, we calculated copy-number gain and loss profiles across clusters (Figure S2d, Table S5). As previously reported^35^, Clusters A and B, corresponding predominantly to Luminal A and Luminal B tumors, showed relatively stable copy-number profiles, with fewer high-frequency arm-level alterations. Cluster C, enriched for HER2-positive tumors, also showed a more restricted copy-number pattern relative to the basal-associated clusters. By contrast, Clusters D and E displayed distinct genomic profiles, supporting their separation as biologically different basal-enriched regions of the landscape.

Cluster E showed the most characteristic basal-like copy-number pattern, including recurrent gains of 1q, 8q, 10p, and 12p, together with losses of 8p and 16q (Figure S2d). By comparison, Cluster D showed a different alteration profile, including enrichment for 4p gain and 5p loss.

Although region F did not form a stable consensus cluster, it exhibited a highly unstable copy-number profile (Figure S2d), consistent with its transcriptionally atypical position outside the main embedding. Region F showed recurrent gains involving 4q, 5q, 8p, 14q, and 16q, together with losses involving 6q, 17p, 18p, 20q, and 22q (Figure S2e).

Together, these data show that the transcriptional clusters are supported by distinct genomic architectures, with the strongest copy-number instability observed in basal-associated regions, particularly Cluster E and region F.

Because mutation calling directly from bulk RNA-seq can introduce false-positive variant calls, we did not infer somatic mutations from the integrated transcriptomic datasets. Instead, we used curated TCGA mutation calls derived from matched DNA sequencing for samples with available data and projected recurrent clinically relevant mutations, including PIK3CA, AKT1, ESR1, ERBB2, BRCA1, BRCA2, and PALB2, onto the PaCMAP landscape (Figure S2f). This analysis enabled visualization of how mutation-defined therapeutic biomarkers distribute across the transcriptional landscape while avoiding artifacts associated with RNA-seq–based mutation detection.

### Differential gene expression analysis reveals distinct biological functions across breast cancer landscape

Leveraging the large sample size of our integrated cohort, we performed differential gene expression analysis to identify transcriptional programs that distinguish and connect breast cancer subtypes. This analysis revealed both subtype-specific pathway enrichments and shared biological programs, highlighting a structured organization of tumor states across the landscape.

Cluster A, comprised primarily of Luminal A tumors exhibited a canonical endocrine program, marked by estrogen receptor signaling and estrogen-dependent gene expression. This was accompanied by lipid metabolism, eicosanoid synthesis, cytochrome P450 activity, and biological oxidation pathways, reflecting coordinated hormone-associated metabolism.

Additional enrichment of extracellular matrix degradation and glycosylation pathways suggests active but regulated microenvironmental remodeling (Table S6-7, Figure 3a).

**Figure 3.**
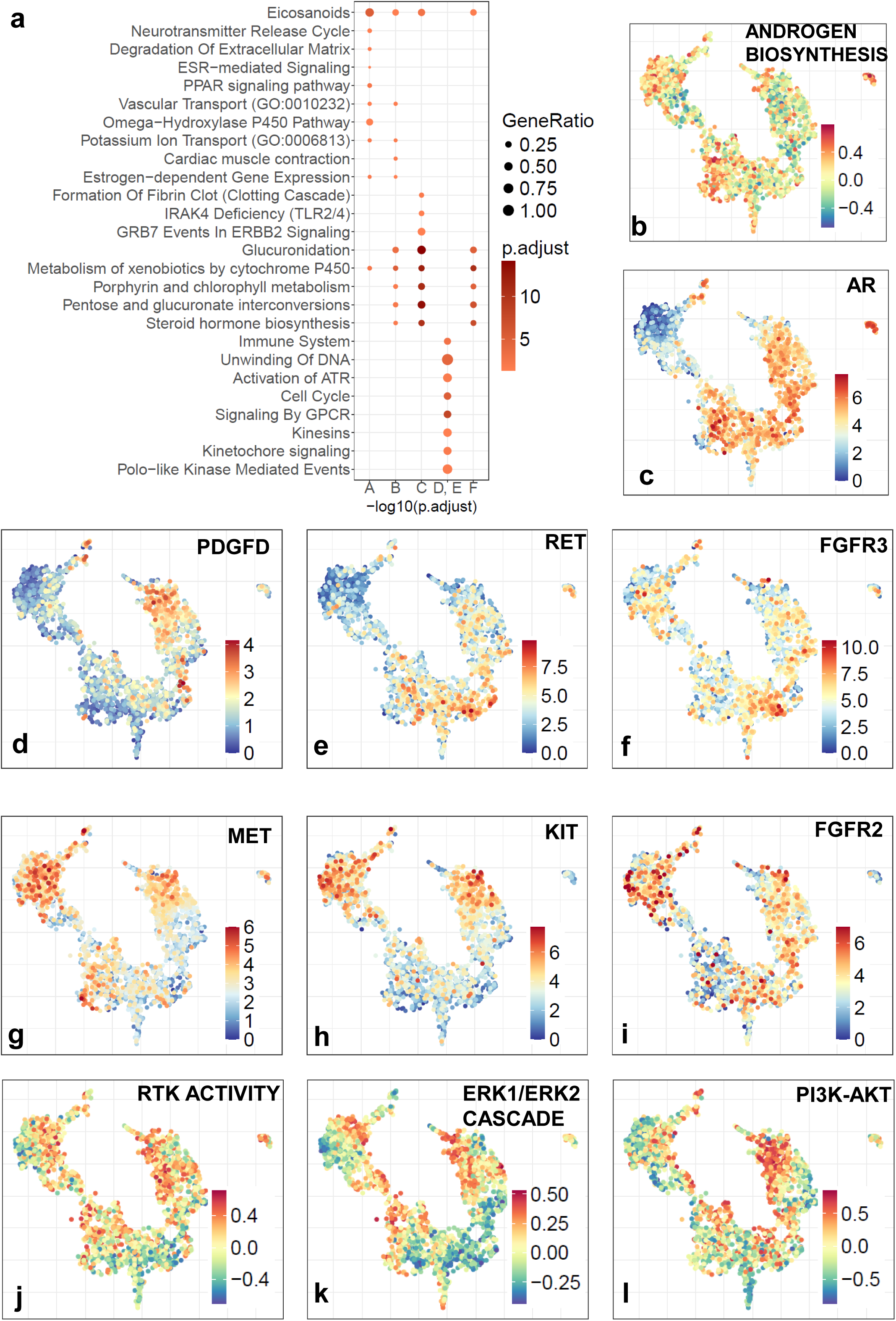
Subtype-specific pathway programs reveal distinct biological axes. (A) Dot plot of differentially expressed pathways across subtypes, highlighting subtype-specific and shared transcriptional programs. (B-C) PaCMAP colored in by GSVA scores for Androgen Biosynthesis pathway and AR gene expression respectively. (D–I) Selected kinases enriched in HER2, Luminal A, Luminal B, and normal-like subtypes, respectively. (J-L) GSVA scores for Tyrosine Kinase receptor signaling, ERK1/ERK2 and PI3K-AKT signaling.

Cluster B retained the core endocrine program but showed glucuronidation and steroid biosynthesis pathways, suggesting altered hormone metabolism. Because glucuronidation can regulate estrogen signaling and contribute to endocrine resistance, these features, together with receptor-mediated signaling, neurotransmitter/post-synaptic pathways, and CYP2E1-related activity, support a more signaling-active and potentially endocrine-resistant Luminal B state (Table S6-7, Fig 3a).

Cluster C, which was predominantly HER2 tumors, was characterized by coordinated ERBB2 signaling coupled with extensive metabolic rewiring. In addition to canonical pathways such as GRB7-mediated ERBB2 signaling (Fig 3a) and integrin–MAPK linkage, these tumors showed enrichment of xenobiotic metabolism, phase I/II conjugation, lipid and fatty acid metabolism, and eicosanoid synthesis. Enrichment of heme/porphyrin metabolism, bile acid processing, complement cascade, TLR signaling, and coagulation pathways further indicates a metabolically plastic and microenvironmentally interactive phenotype. Notably, Cluster C also contained the largest number of metastatic samples from the CMI-MBC cohort, including four of the five liver metastases, raising the possibility that metastatic site composition may contribute to, or reflect, some of the metabolic features enriched in this region (Table S6-7).

Although Clusters D and E were both enriched for basal-like tumors, they separated into distinct transcriptional programs. Cluster E showed the strongest proliferation-associated signature, with enrichment of cell cycle, mitotic checkpoint, sister chromatid cohesion, chromatid separation, and Polo-like kinase pathways, consistent with a highly proliferative basal state. In contrast, Cluster D showed a more immune- and epithelial-associated basal program, with enrichment of keratinization, cornified envelope formation, antimicrobial peptides, and metal sequestration pathways, suggesting a barrier-like and immune-reactive phenotype (Figure 3a).

Beyond these differences, both basal-associated clusters showed evidence of broader signaling and structural remodeling. Enrichment of GPCR ligand binding, rhodopsin-like receptor pathways, extracellular matrix organization, collagen biosynthesis, and developmental signaling, including FGFR2-related pathways, suggests enhanced microenvironmental sensing, matrix interaction, and signaling plasticity across basal tumors (Figure 3a; Tables S6–7).

Notably, cluster F was clinically annotated as triple-negative due to lack of ER, PR, HER2 status, and showed an AR-enriched and luminal-like transcriptional profile rather than a canonical basal-like state. This region showed high expression of AR, FOXA1, GATA3, AGR2, KRT8, KRT18, KRT19, MUC1, SPDEF, TFF3, SCUBE2, CA12, and low expression of basal markers including KRT5, KRT14, SOX10, FOXC1, and TP63 (Figure S2a, Figure 2). Despite low ESR1 and PGR, region F retained luminal-regulatory and epithelial features and showed partial HER2-adjacent expression, including variably elevated ERBB2 and PGAP3. These findings support region F as an atypical AR-enriched, HER2-like TNBC region distinct from the immune-enriched and proliferative basal-associated clusters.

### Kinase and Receptor tyrosine kinase programs distinguish signaling states across landscape clusters

We identified numerous kinase-associated genes enriched across subtype-defined regions of the landscape (Table S8). The expression of many kinase genes was strongly cluster-specific.

Cluster A, corresponding predominantly to Luminal A tumors, was enriched for kinases, their ligands, and kinase-associated genes linked to stromal interaction, metabolic regulation, and differentiated receptor signaling, including PDGFD (Figure 3d), PDK4, TIE1, TGFBR2 (Figure S3a), BMX, NTRK1, and PRKG1 (Table S8), in comparison to cluster D and E. This profile is consistent with a luminal-associated state shaped by endocrine and stromal signaling rather than overt proliferative kinase activation.

Cluster B, corresponding predominantly to Luminal B tumors, showed increased expression of kinases such as RET^36^ and FGFR3^37^ (Figure 3e-f), along with FGFR1, IGF1R, and WNK4 (Figure S3a), supporting a more signaling-active luminal state. Additionally, enrichment of HSPB8^38^, previously linked to poor prognosis and tamoxifen resistance, and MPP7, a potential prognostic biomarker and therapeutic target for clear cell renal cell carcinoma (Figure S3a) further suggests that Cluster B contains kinase-linked stress-response and survival programs that may contribute to its more aggressive luminal phenotype.

Cluster C, enriched for HER2-positive tumors, showed the expected activation of HER2-associated signaling, including high ERBB2 expression (Figure 2a), together with MPP7^39^, AKT1^40^ (Figure S3a), and PIP5K1B^41^, in comparison to cluster D and E. This pattern is consistent with growth factor receptor signaling coupled to MAPK, PI3K–AKT, and phosphoinositide pathway activity.

The basal-associated Clusters D and E showed the broadest kinase enrichment in relation to the other clusters. These clusters showed elevated expression of receptor tyrosine kinases and signaling mediators including MET^42^, KIT^43^, and FGFR2^37^ (Figure 3g–i), suggesting activation of growth factor, invasive, and developmental signaling programs.

Consistent with this pattern, Cluster D and a subset of Cluster A showed elevated GSVA scores for receptor tyrosine kinase activity, ERK1/ERK2 cascade, and PI3K–AKT signaling, indicating that growth factor–linked signaling programs extend beyond canonical HER2-enriched regions and are active in selected basal-associated and luminal-associated states (Figure 3 j-l). Basal-associated clusters also showed elevated expression of YES1 and PIM1 (Table S8), both previously proposed as therapeutic targets in TNBC, as well as TTK^44^ (Figure S3a), a mitotic checkpoint kinase linked to proliferation and survival.

Together, these results show that kinase enrichment reflects distinct pathway architectures across the landscape: stromal/metabolic signaling in Cluster A, receptor and survival signaling in Cluster B, canonical HER2-growth factor signaling in Cluster C, and broad RTK, MAPK/PI3K, and mitotic kinase activation in basal-associated Clusters D and E. These cluster-specific kinase programs provide candidate signaling vulnerabilities for downstream therapeutic interrogation.

### Basal-associated clusters (cluster D and E) resolve into immune-enriched and proliferative states

The basal-associated regions of the landscape showed two distinct biological programs. Cluster E was marked by high GSVA scores for cell cycle and DNA replication pathways (Figure 4a–c), consistent with a strongly proliferative basal state. By contrast, Cluster D showed evidence of an immune-enriched and lower-purity microenvironment. The T cell–inflamed gene expression profile (GEP)^45^, which captures activated T cell and interferon signaling, was elevated in Cluster D (Figure 4d), supporting the presence of an immune-active tumor microenvironment.

**Figure 4.**
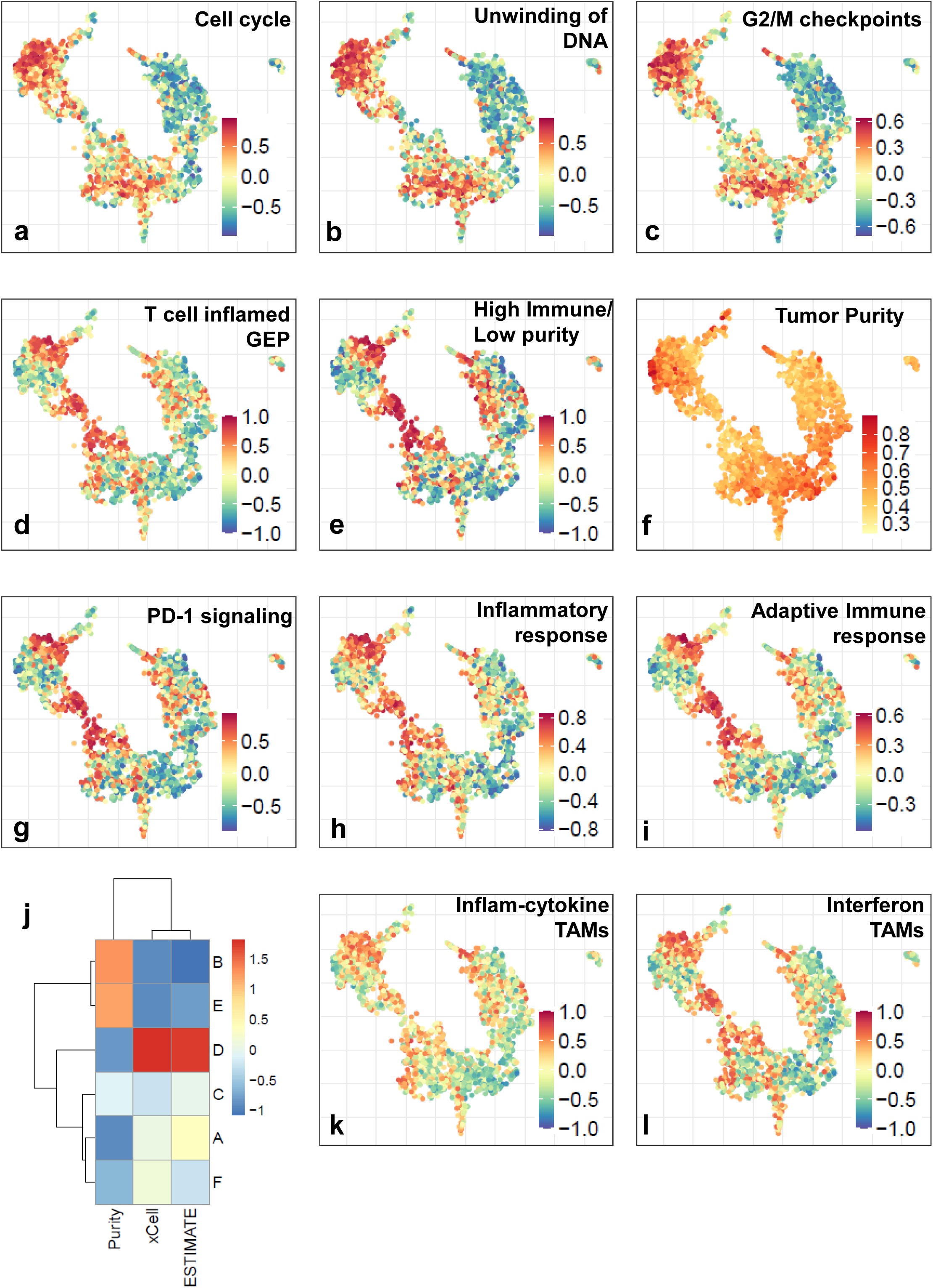
Basal tumors are characterized by proliferative and immune-enriched programs. (A–C) GSVA scores for cell cycle and DNA replication pathways, showing strong enrichment in basal regions. (D) T cell–inflamed gene expression profile (GEP) enriched in basal tumors. (E) Immune gene signature showing high immune scores and low tumor purity in basal regions. (F) Tumor purity projected onto the landscape, with basal regions exhibiting lower purity. (G-I) GSVA scores for PD-1 signaling, Inflammatory response and Adaptive immune response. (J) Heatmap of median scores for ESTIMATE, xCell immune scores and tumor purity scores in each cluster. (K–L) TAM-associated gene signatures projected onto the landscape, including inflammatory cytokine TAMs, interferon-responsive TAMs, immune-regulatory TAMs, Kupffer-like TAMs, MT-RTM macrophages

This pattern was further supported by an independent five-gene immune signature^46^, which showed high immune scores and low tumor purity in Cluster D (Figure 4e). Direct estimates of tumor purity across the landscape calculated using PUREE^47^, confirmed lower tumor purity in the Cluster D and a subset of cluster A tumors (Figure 4f), suggesting increased contribution from non-tumor components, including immune and stromal cells. In line with this interpretation, PD-1 signaling, inflammatory response programs (Fig 4g-h), CD8 T cells, regulatory T cells, M1 macrophages, and dendritic cell signatures were enriched in Cluster D (Figure S4a), accompanied by elevated expression of immune checkpoint genes PDCD1, CD274, LAG3, CTLA4, and HAVCR2/TIM3 (Figure S4b). These findings indicate coordinated immune infiltration and checkpoint regulatory activity within Cluster D. Consistent with these findings, orthogonal immune-deconvolution approaches, including ESTIMATE, xCell, and CIBERSORTx, converged on the same pattern: Cluster D showed the strongest immune enrichment and the lowest inferred tumor purity across the landscape (Fig. 4j; Fig. S4c).

We also evaluated the postpartum breast cancer (PPBC)–associated signature reported by Jindal et al.^48^ (Fig. S4c), derived from breast cancer in young women diagnosed postpartum versus nulliparous controls. Although originally defined in ER-positive disease, this cell-cycle and T cell activation-associated program was most enriched in Clusters D and E in our landscape, suggesting that it captures an aggressive proliferative and immune-active state aligned more closely with basal-associated biology than with classic hormone replacement programs.

Together, these findings indicate that the two basal-associated clusters represent distinct but related tumor states: Cluster E is dominated by proliferation and DNA replication, whereas Cluster D is characterized by immune enrichment, reduced tumor purity, and inflammatory microenvironmental activation.

### Tumor microenvironmental cell contributions are regionalized across the landscape

Because bulk RNA-seq captures aggregate expression from tumor cells and the surrounding microenvironment, we used curated gene signatures to evaluate whether non-tumor cell-associated programs varied across landscape regions. We found that several non-tumor cell types were associated with specific transcriptomic tumor subtypes.

#### Tumor-associated macrophage programs distinguish Cluster D from Cluster E within basal-associated regions

Tumor-associated macrophages (TAMs) are a major component of the tumor microenvironment and play critical roles in regulating inflammation, immune suppression, tissue remodeling, and therapeutic response^49,50^. Distinct TAM states have been described across cancers, reflecting functional specialization ranging from pro-inflammatory and interferon-responsive phenotypes to tissue-resident and immunoregulatory programs^51^.

GSVA-based projection of TAM gene signatures derived from published studies^52^ (Table S2) revealed strong cluster-specific enrichment in Cluster D. This region showed marked enrichment of inflammatory cytokine-associated TAMs, as well as interferon-responsive TAM programs (Figure 4k-l), consistent with an immune-activated microenvironment. Immune-regulatory TAM signatures were also elevated in Cluster D (Figure S4d), suggesting concurrent activation of immunosuppressive macrophage programs. Tissue-resident macrophage signatures, including Kupffer cell–like TAMs and MT-RTM macrophages, localized to distinct regions of the landscape (Figure S4d), indicating structured heterogeneity in macrophage states.

Notably, FOLR2+ TAMs, associated with tissue-resident and homeostatic macrophage phenotypes in breast cancer^53^, were enriched in Cluster D and in a subset of the Cluster A/Luminal A region (Figure 4l), suggesting that tissue-resident macrophage programs are shared between the immune-enriched basal-associated region and selected luminal-associated tumors.

Together, these findings demonstrate that various tumor associated macrophages are not uniformly distributed but instead map to distinct regions of the transcriptional landscape. Cluster D showed the strongest enrichment of inflammatory, interferon-associated, and immune-regulatory macrophage programs, consistent with its enrichment for the immune-modulatory^26^ TNBC subtype in the subtype-composition analysis (110/181 samples, 60.8%; Fig. S2b). highlighting the complexity of its immune microenvironment. These results suggest potential relevance for immunomodulatory therapeutic strategies.

#### Cancer-associated fibroblast programs further stratify cluster-specific tumor microenvironments

Cancer-associated fibroblasts (CAFs) are key regulators of the tumor microenvironment, contributing to extracellular matrix remodeling, immune modulation, angiogenesis, and metabolic support^52,54,55^. To assess stromal heterogeneity across the landscape, we quantified CAF-associated gene signatures curated from published studies using GSVA (Table S2).

CAF programs showed clear cluster-specific enrichment (Figure 5a–g). Progenitor CAF and myCAF signatures were most prominent in Cluster A, while ECM CAF was enriched primarily in Cluster A and secondarily in Cluster C, consistent with matrix-remodeling activity in luminal- and HER2-associated regions. Notably, apCAF enrichment in Cluster D paralleled the immune-enriched and immune-modulatory features of this region, whereas metabolic CAF enrichment in Cluster E aligned with its highly proliferative basal-associated state and enrichment for the BL1 TNBC program (161/236 samples, 68.2%; Fig. S2b). Together with the TAM analyses, these findings indicate that distinct immune and stromal programs are coordinated with tumor transcriptional state across the landscape.

**Figure 5.**
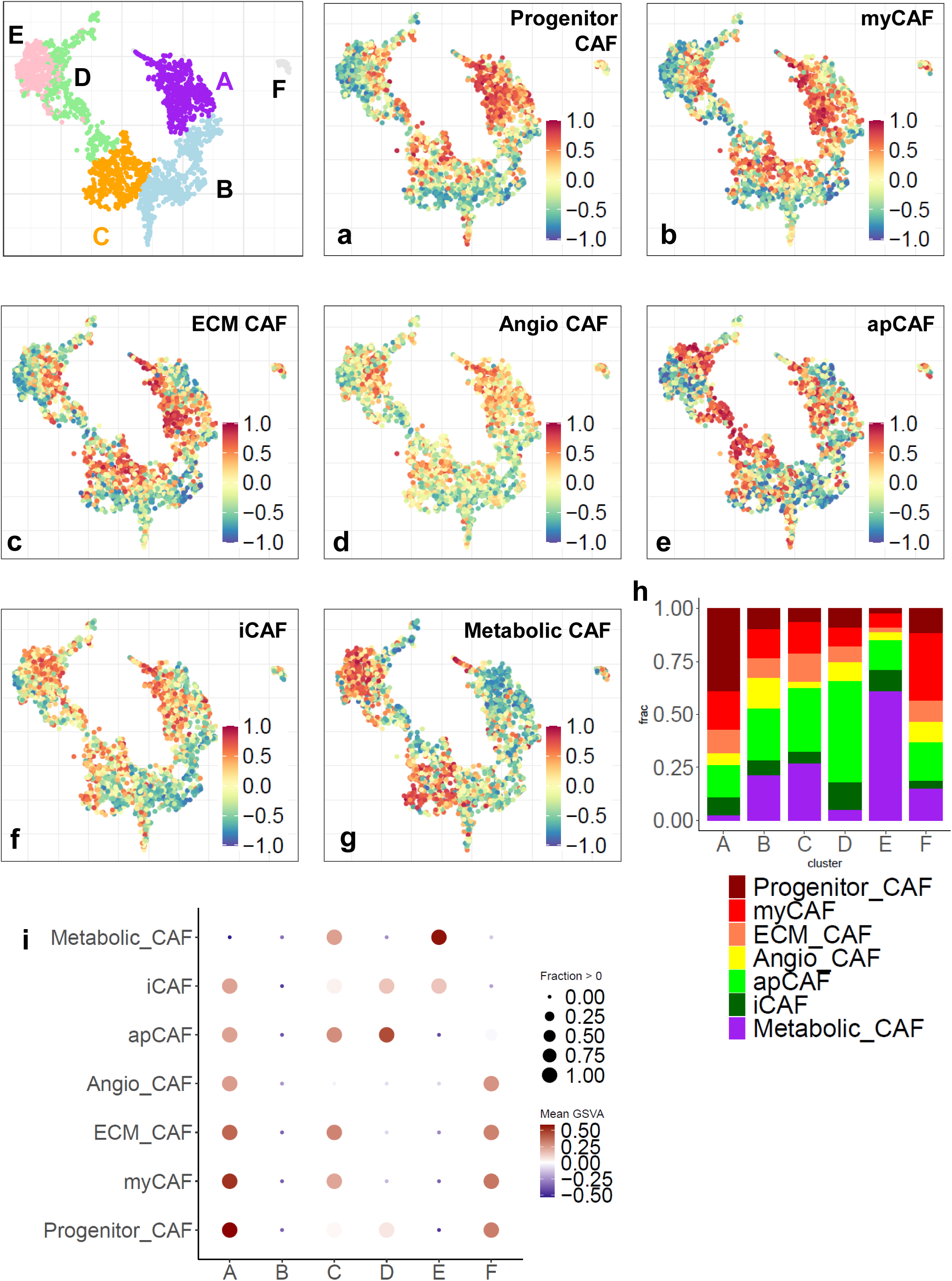
Basal subtype heterogeneity and stromal programs define microenvironmental structure. (A–G) CAF-associated gene signatures projected onto the landscape, including progenitor CAF, myCAF, ECM CAF, angiogenic CAF, apCAF, iCAF and metabolic CAF programs, revealing distinct stromal distributions. (H) Stacked bar plot showing the fraction of samples in each landscape cluster assigned to each dominant CAF program, defined as the CAF signature with the highest GSVA score in that sample (I) Dot plot summarizing CAF program enrichment across landscape clusters. Dot color represents the mean GSVA score for each CAF signature within each cluster, and dot size represents the fraction of samples with positive enrichment score greater than zero.

We summarized these patterns using both dominant CAF-state assignment and a cluster-by-CAF dot plot (Figure 5h,i). These analyses confirmed that CAF programs are not uniformly distributed but instead form cluster-specific stromal compositions. Representative marker genes for apCAF, metabolic CAF, progenitor CAF, and ECM CAF localized to the corresponding regions of the landscape (Figure S5a–d), further supporting the GSVA-based CAF-state assignments. Together, these findings identify distinct CAF-associated transcriptional programs aligned with tumor transcriptional state.

#### Neuron content and synaptic signaling programs are enriched in basal-associated tumor subtypes

A subset of breast cancers is known to be innervated^56^, and nerve fibers within the tumor microenvironment have been associated with nerve growth factor production, lymph node invasion, and aggressive disease features. In addition to kinases, we also observed a number of synaptic genes that were differentially regulated across the clusters (Table S9). Genes associated with synaptic signaling and neuronal communication have increasingly been recognized as contributors to tumor cell–cell interaction, vesicle trafficking, and receptor-mediated signaling in cancer. These programs are thought to reflect co-option of neuronal-like signaling modules that enhance tumor plasticity, communication, and microenvironmental sensing.

Clusters D and E, the two basal-associated regions, stand out among the breast-cancer subtypes with the most consistent expression of many neuron-specific genes across known programs.

These clusters show the highest level of expression for transcription factors involved in neuronal development and differentiation including POU4F1, HAND2, and SOX11 (Fig 6a-c), which drive expression of other neuronal genes. They express UCHL1 (Fig 6d), a specific marker of neurons of the spinal cord and peripheral nervous system. They also express receptors for cholinergic signaling with high levels of CHRNA5, CHRM1, and CHRM3(Supp Fig 6a-c).

**Figure 6.**
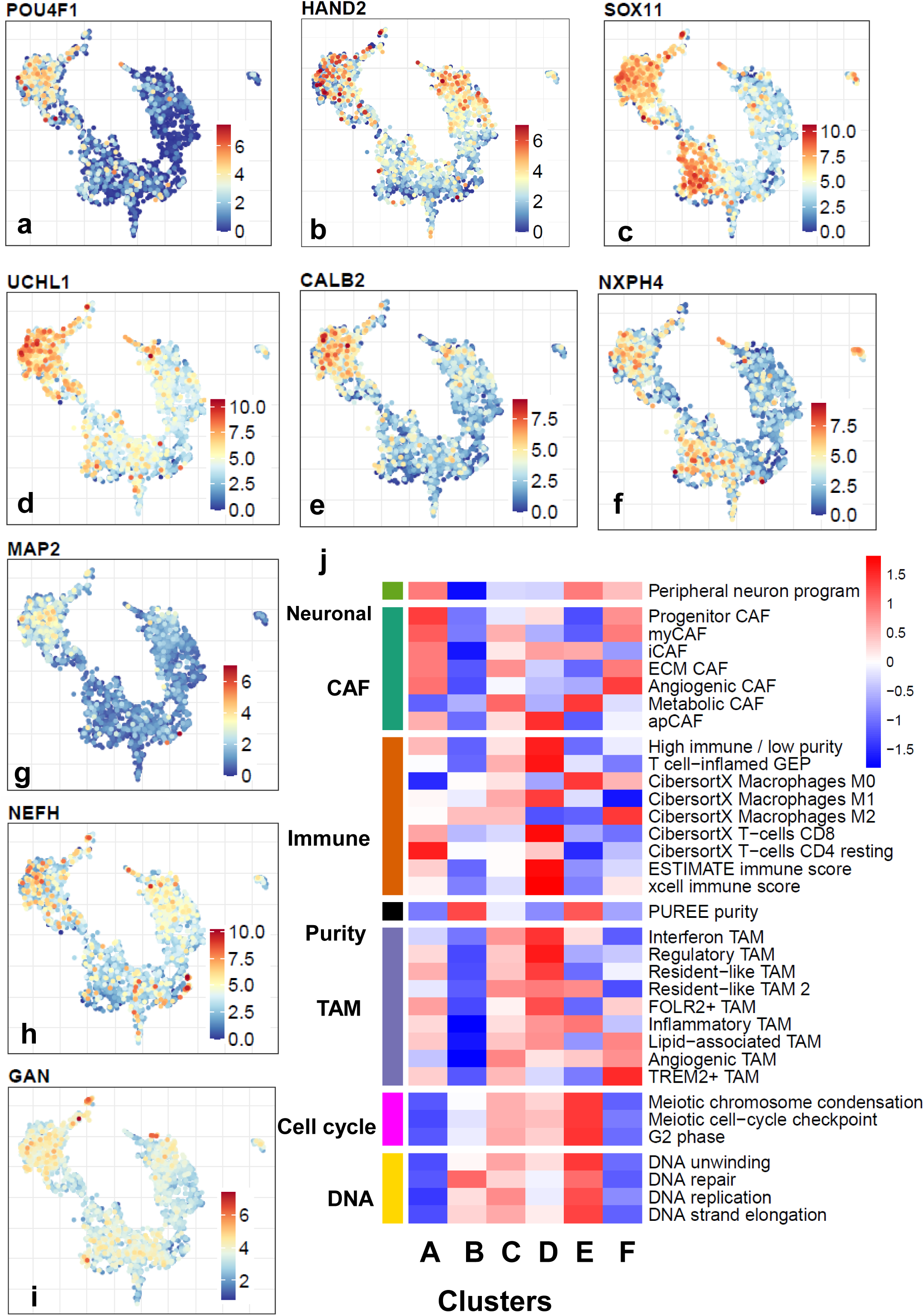
Neuronal-like and synaptic gene expression across the breast cancer landscape. **(A–I)** PaCMAP embedding colored by expression of selected neuronal-like and synaptic-associated genes. (A) POU4F1, (B) HAND2, (C) SOX11, (D) UCHL1, (E) CALB2, (F) NXPH4, (G) MAP2, (H) NEFH, and (I) GAN show enrichment across basal-associated and selected luminal-associated regions, supporting shared expression of neuronal filament and microtubule-associated programs across multiple clusters. (J) Heatmap of mean GSVA scores for cell cycle, DNA replication, immune deconvolution programs, purity scores, TAM, and CAF programs across clusters, summarizing coordinated tumor-intrinsic and microenvironmental programs.

Other unique neuronal markers of basal tumors include the sympathetic secondary neuropeptide GAL, CALB2 (Fig 6e) involved in calcium ion buffering within neurons, and NXPH4 (Fig 6f) a neurosecretory glycoprotein. Although expression of these genes is unique, basal-associated regions most consistently share expression of neuronal markers with other breast cancer subtypes.

Several neuronal programs were shared between basal-associated clusters and Cluster A, the Luminal A–enriched region. These included markers of neuronal filaments and microtubules including NEFH, MAP2, and GAN. They together express receptors for sympathetic autonomic signaling including ADRA2A, ADRA2B, ADRB2, and NPY1R. They also express MBP and S100B, markers of myelination and Schwann cell infiltration at the highest level. Taken together these two subtypes trend as the most neuron-like in their transcriptome, and potentially the most interactive with the peripheral nervous system.

Cluster A tumors also expressed receptors for neurotrophic factors including NGFR, NTRK2, and GFRA1. They uniquely express TAC1 and TACR1, the ligand-receptor pair for inflammatory nociceptive Substance P signaling. Luminal A tumors most consistently express components of voltage gated ion channels for sodium, potassium, and calcium including SCN2B, SCN4A, SCN4B, KCNA3, CACNA1C and CACNA1D. Luminal A also expresses oxytocin receptor OXTR at the highest level, Post synaptic marker SHANK3, as well as the immediate early gene and protooncogene FOS. These implicate Luminal A tumors as the most excitable of the subtypes.

Cluster B, the Luminal B–enriched region, was distinguished by enrichment of presynaptic and vesicle-associated machinery. This included SYP, SNAP25, SYN1, VAMP2, STX1A, CLTC, and SV2A, along with postsynaptic scaffolding genes DLG3 and DLG4. These patterns suggest that Cluster B is enriched for synaptic vesicle trafficking and membrane-fusion programs, consistent with a signaling-active luminal state. It is also high in MAPT with luminal A tumors, essential for generating tau, the neuronal cytoskeletal protein whose aggregation is associated with Alzheimer’s disease. These implicate Luminal B tumors as the most-rich in synaptic machinery.

Cluster C, enriched for HER2-positive tumors, did not show a single dominant neuronal program but shared several synaptic-associated genes with other clusters, including SOX11, NXPH4, GAN, STX1A, CHRNA1, CHRM1, KCNA3, and CLTC. This suggests that these tumors may utilize neuronal machinery from a wide array of processes without forming a uniquely synaptic transcriptional state.

Together, these findings suggest that synaptic and neuronal-associated gene expression profiles are not uniformly distributed across the landscape. Instead, basal-associated clusters show the broadest neuronal-like program, Cluster A is enriched for neurotrophic and excitability-associated features, Cluster B is enriched for synaptic vesicle machinery, and Cluster C displays a more heterogeneous pattern. These programs may represent additional communication and signaling axes that contribute to cluster-specific tumor behavior and microenvironmental interaction.

#### CAF, TAM, and neuronal-like programs define cluster-specific tumor states

To summarize the relationships between tumor-intrinsic and microenvironmental tumor programs, we computed mean GSVA scores for proliferative, immune, TAM, CAF, and synaptic signaling signatures across clusters and visualized them in an integrated heatmap (Figure 6j).

This analysis reinforced the distinct biology of the cluster-defined regions. Cluster D was characterized by high immune, TAM, and checkpoint-associated scores together with low tumor purity, consistent with an immune-enriched basal-associated state. By contrast, Cluster E showed the strongest enrichment of cell cycle and DNA replication programs, defining a highly proliferative basal-associated state. Cluster A was enriched for stromal and extracellular matrix– associated CAF programs together with selected synaptic and neurotrophic signatures, whereas Cluster B showed a more signaling-active luminal profile but was relatively depleted for CAF, immune, and TAM-associated programs, suggesting a tumor-intrinsic luminal state with limited microenvironmental contribution. Cluster C displayed intermediate stromal and signaling features, consistent with its HER2-associated identity. Together, these data demonstrate that proliferative, immune, stromal, and neuronal programs are tightly coordinated and jointly define distinct regions of the transcriptional landscape.

### Transcriptional landscape predicts potential regional organization of therapeutic sensitivity and resistance

Patients who meet the same clinical criteria for a therapy can differ substantially in response because tumors within a treatment-defined group may still have distinct underlying biology. Some patients achieve sustained benefit, whereas others show primary resistance or relapse after an initial response ^2,3^. We therefore asked whether our landscape could be used to predict drug response and resistance programs according to tumor biology rather than clinical indication alone.

We first examined whether treatment annotations were available across the integrated cohort. However, treatment metadata were available for fewer than 1% of samples, limiting their utility for landscape-wide analysis. When hormone therapies were considered collectively, treated samples localized primarily to Clusters A and B, consistent with their Luminal A and Luminal B composition, whereas radiotherapy annotations did not show a clear spatial pattern (Figure S7a,b).

Because these sparse annotations were insufficient to evaluate response or resistance across the landscape, we instead curated published studies^57–59^ with matched responders and non-responders and used genes upregulated in non-responders (Table S2) to generate GSVA-based resistance scores. Resistance signatures were generated using three therapies based on currently available datasets, although the approach is readily expandable to additional therapies as relevant data become available.

For HER2-positive disease, Trastuzumab (Herceptin) is a standard treatment. Trastuzumab is a monoclonal antibody that targets ERBB2/HER2, thereby inhibiting HER2-driven signaling and tumor growth. Figure 7a,b shows the distribution of HER2-positive tumors and ERBB2 expression across the landscape. HER2 protein expression, available for the TCGA subset, was concordant with the observed ERBB2 RNA expression pattern. Using a chemo-resistance signature derived from Dadiani et al^57^, who identified 122 genes upregulated in nonresponders, we observed low predicted resistance scores within the HER2-enriched region, whereas Luminal A, Luminal B, and basal regions showed higher predicted resistance to trastuzumab (Figure 7c).

**Figure 7.**
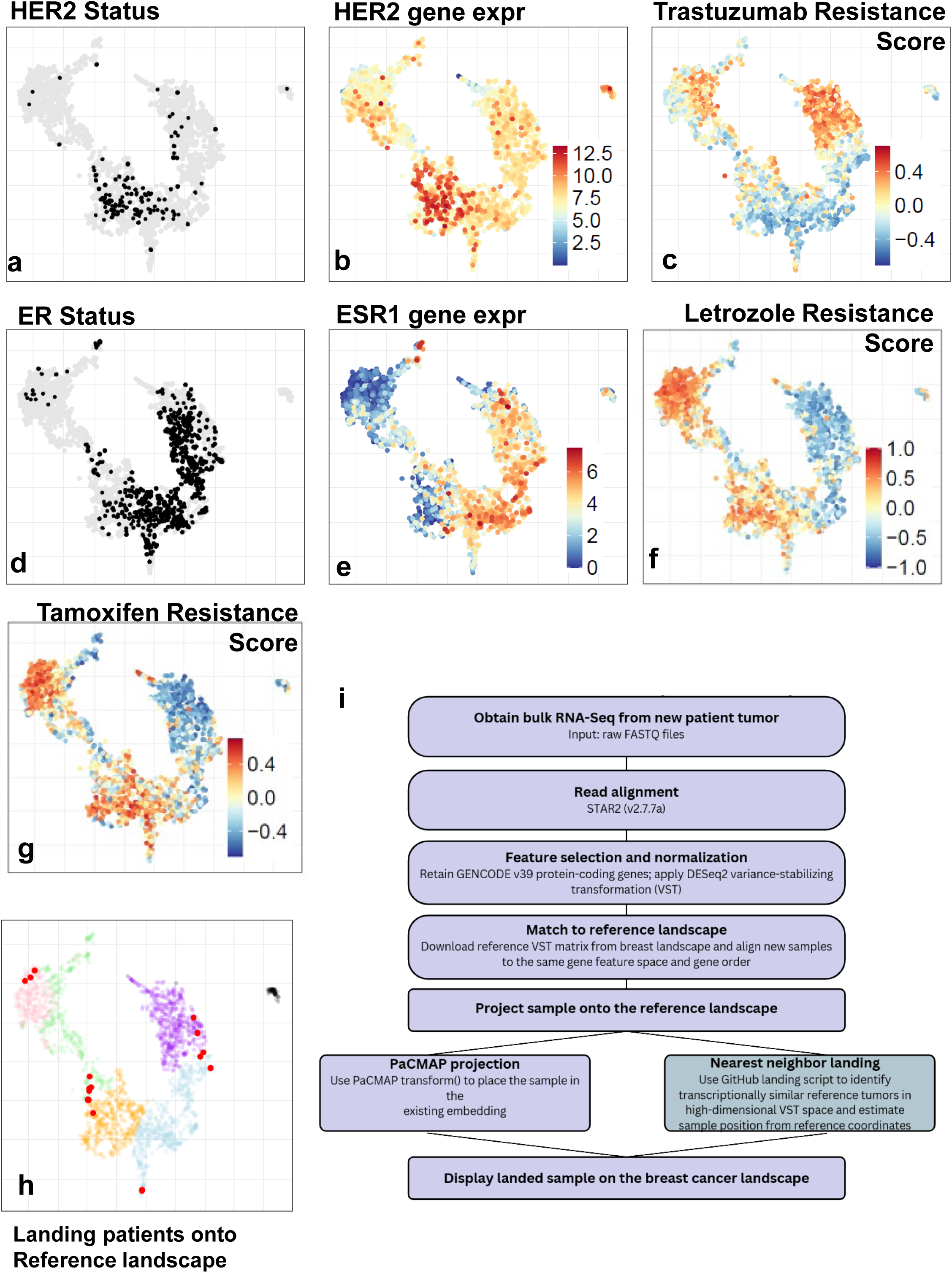
Therapeutic response and resistance programs map to distinct regions of the breast cancer landscape. (A) PaCMAP embedding colored by reported HER2 status, showing localization of HER2-positive tumors. (B) Gene expression of ERBB2 projected onto the landscape. (C) GSVA-based Trastuzumab resistance score projected onto the landscape, with blue indicating predicted responsiveness and red indicating predicted resistance. (D) PaCMAP embedding colored by reported ER-positive status, showing localization of hormone receptor–positive tumors. (E) Gene expression of ESR1 projected onto the landscape. (F-H) GSVA-based Letrozole and Tamoxifenresponse/resistance score projected onto the landscape respectively, with blue indicating predicted responsiveness and red indicating predicted resistance. (H) Projection of 19 independent patient samples from GSE254461 onto the breast cancer reference landscape, showing that external tumors land within subtype-consistent regions of the embedding. (I) Schematic of the patient-landing workflow. External samples were processed to match the reference expression matrix and projected onto the landscape using both PaCMAP transform and a nearest-neighbor landing approach, enabling comparison of projected sample position with reference cluster and PAM50 subtype structure.

For hormone receptor–positive tumors, aromatase inhibitors, such as letrozole (Femara), are widely used. Letrozole suppresses estrogen production by inhibiting aromatase, thereby reducing ER-driven transcriptional signaling. Figure 7e-g shows the distribution of hormone receptor– positive tumors, ESR1 expression and estrogen receptor signaling. Using a nonresponse signature derived from Lee et al^58^, who studied 58 ER-positive tumors, we found that Luminal A and Luminal B regions showed lower predicted resistance scores, while HER2-enriched and basal regions showed higher predicted resistance to letrozole (Figure 7h).

Tamoxifen, another commonly used endocrine therapy for HR+ breast cancer, acts as a selective estrogen receptor modulator (SERM), directly antagonizing ER signaling in breast cancer cells. Using gene expression programs associated with tamoxifen-resistant clusters identified by Kim et al^59^ from single-cell analysis of tumors from 10 women with ER-positive/HER2-negative breast carcinoma, we generated a tamoxifen resistance score and projected it onto the landscape. Luminal A and Luminal B regions showed lower predicted resistance scores, whereas HER2-enriched and basal regions showed higher predicted resistance (Figure 7i).

Together, these analyses show that expression profiles correlate with drug response and localize to specific regions of the transcriptional landscape and largely align with expected disease biology. More importantly, they indicate that resistance is not fully explained by receptor status or subtype label alone. Rather, predicted therapeutic response appears to be shaped by the broader transcriptional context in which the targetable marker is expressed.

### Landing external patient samples onto the breast cancer reference landscape

To test whether the landscape could serve as a reusable reference for new tumors, we projected 19 independent patient breast tumor samples from GSE254461 onto the breast cancer landscape (Fig. 7h, Table S10). Samples were landed using two complementary approaches: the open-source PaCMAP transform function and a laboratory-developed nearest-neighbor projection method, summarized in the landing workflow schematic (Fig. 7i). The two methods produced concordant placements for each patient (Fig. S7c). PAM50 assignments for the projected samples were consistent with their landing positions, with basal, HER2-enriched, Luminal A, and Luminal B tumors mapping to the corresponding regions of the reference landscape. Together, these findings demonstrate that the landscape can be used to position external patient samples within established breast cancer transcriptional states and their nearest neighbors can be used to predict their biology.

## DISCUSSION

In this study, we construct a unified and interactive transcriptional landscape of breast cancer by integrating 13 independent transcriptomic datasets. This framework organizes tumors along continuous biological axes and recapitulates canonical subtypes while resolving finer-scale heterogeneity. Unlike traditional classification schemes that rely on a small subset of genes, this approach uses 18,089 protein coding genes to stratify patients and enables visualization of tumor states within a continuous space, providing a more nuanced representation of breast cancer biology.

This landscape provides the ability to interrogate multiple layers of tumor biology within a single framework. By projecting gene expression, pathway activity, kinase programs, and microenvironmental features—including tumor-associated neurons, macrophages and fibroblasts—we identify coordinated biological modules that define distinct regions of the landscape. This integrative view reveals how tumor-intrinsic programs and microenvironmental states are regionally organized and interact across subtypes in the landscape.

The subtype-specific enrichment of kinase programs and synaptic signaling genes highlights additional layers of regulatory heterogeneity across breast cancer. Distinct kinase repertoires observed across subtypes include both well-established oncogenic drivers and regulators with tumor suppressive functions, underscoring the diversity of signaling dependencies that may be therapeutically exploitable. In parallel, the enrichment of synaptic and neuronal signaling genes—particularly in basal tumors—suggests co-option of communication and signaling modules that may contribute to tumor plasticity and microenvironmental interaction. Notably, several of these kinases and synaptic-associated genes are either established drug targets or represent emerging candidates for therapeutic intervention, supporting their potential relevance in guiding subtype-specific treatment strategies.

Importantly, mapping clinical outcomes onto the landscape identifies regions associated with differential survival that extend beyond canonical subtype boundaries. This suggests that transcriptional state, rather than subtype label alone, may better capture clinically relevant variation. In addition, as shown with other transcriptomic landscapes^9,11,12^, one can also project a new patient onto a reference landscape, and from the nearest neighbors of that landing point, predict tumor biology, outcome and therapeutic response.

Finally, the landscape offers a way to examine a common clinical problem: patients may meet the standard criteria for a given therapy yet their disease fails to respond, or relapses after an initial response, because the broader transcriptional biology of their tumor differs from what is implied by receptor status alone. To test this concept, we mapped therapy-associated response and resistance signatures onto the landscape as an initial proof of concept. We focused on therapies for which published responder and nonresponder expression signatures were available, including the HER2-targeted agent trastuzumab and endocrine therapies such as tamoxifen and letrozole. Projection of these signatures onto the landscape identified regions enriched for predicted sensitivity or resistance and suggested that therapeutic response aligns with continuous tumor states rather than discrete subtype labels alone. However, this analysis is not intended to represent the full treatment landscape of breast cancer. Clinically important therapies, including PARP inhibitors, PI3K-pathway inhibitors, immune checkpoint inhibitors, selective estrogen receptor degraders, antibody–drug conjugates, and additional targeted agents, will require incorporation of matched pretreatment transcriptomic and response data. As such datasets become available, this framework can be useful for mapping treatment sensitivity and resistance across breast cancer states.

Together, these findings establish the landscape as a reusable and interactive resource for exploring tumor heterogeneity, linking molecular programs to clinical outcomes, and identifying potential therapeutic vulnerabilities.

### Limitations of the study

Several limitations should be considered when interpreting our results. First, the landscape is constructed from bulk transcriptomic data and thus represents aggregated signals from both tumor cells and the surrounding microenvironment. While this enables robust characterization of global transcriptional structure, it does not provide cell-type–specific resolution. Second, integration of datasets generated across multiple studies and sequencing platforms may introduce technical variability. Although we implemented a consistent preprocessing pipeline and batch correction and observed strong intermixing of datasets within the embedding, the possibility of residual confounding cannot be entirely ruled out. Future studies incorporating prospective data, as well as single-cell and spatial transcriptomic approaches, will help refine the resolution and biological interpretation of the landscape. Finally, inference of therapeutic sensitivity based on gene expression signatures is correlative and does not establish causal or functional dependencies, necessitating experimental validation.

## STAR METHODS

### KEY RESOURCES TABLE

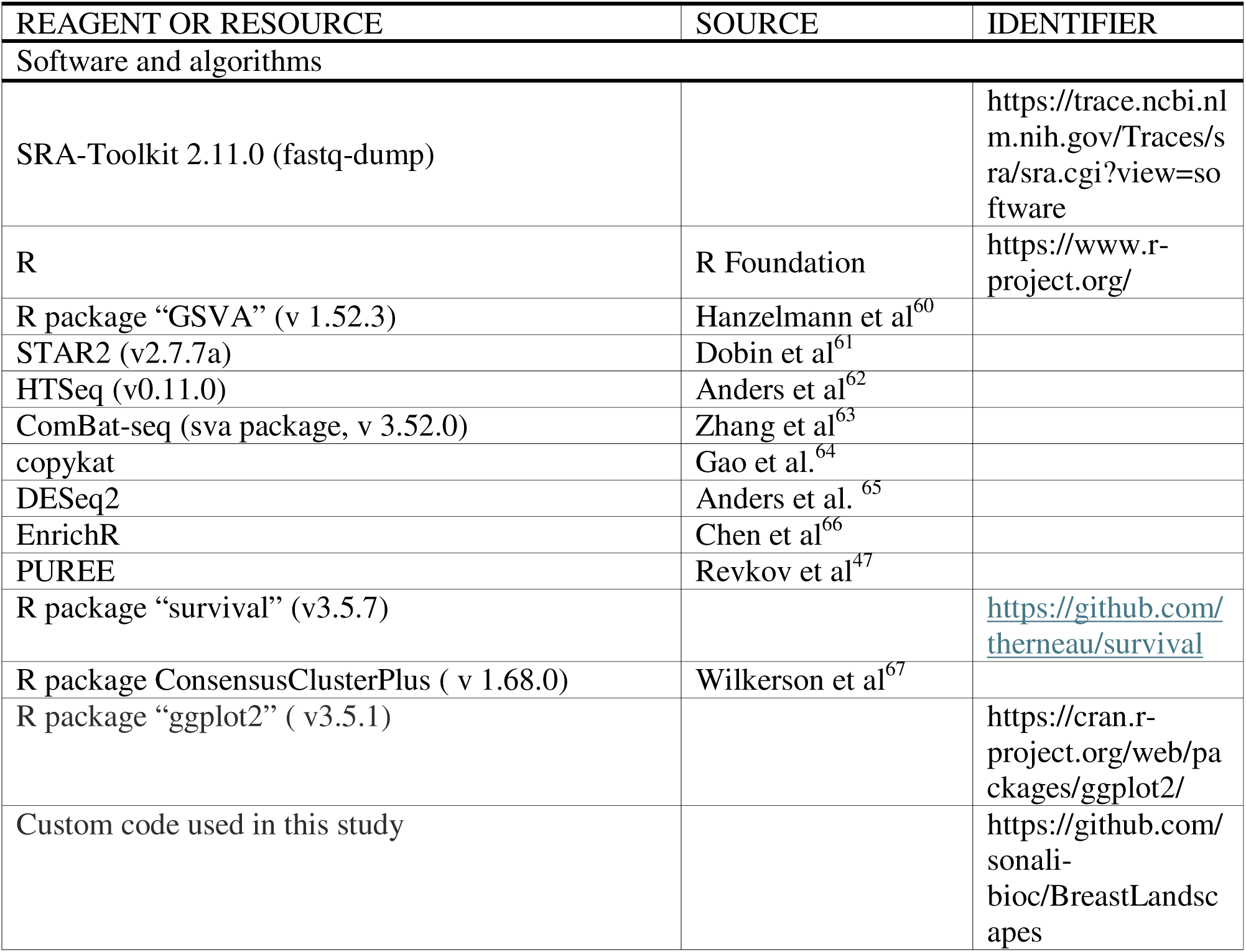

#### Data Collection and Processing

Publicly available bulk RNA sequencing datasets were obtained from GEO and GDC repositories. Raw FASTQ files were uniformly processed to ensure cross-cohort consistency.

Quality control was performed using FastQC (v0.11.9) and MultiQC (v1.9). Reads were aligned to the GENCODE GRCh38.primary_assembly reference genome using STAR (v2.7.7, two-pass mode). Gene-level counts were generated with HTSeq (v0.11.0) using GENCODE v39 annotations.

Count matrices were aggregated across cohorts. Batch effects were corrected using ComBat-seq (sva), and variance-stabilized transformation (VST) values for only protein coding genes, were computed using DESeq2 for downstream visualization and analysis.

#### Dimensionality Reduction and Clustering

Dimensionality reduction was performed using t-SNE, UMAP, and PaCMAP. PaCMAP embeddings were selected as the reference landscape due to optimal preservation of local and global structure. Consensus clustering was applied to relevant subsets to identify stable transcriptional states.

#### Differential Expression and Pathway Analysis

Differential gene expression analysis was performed using DESeq2, with significance defined as FDR < 0.05 and |log fold change| > 0.5 pairwise between all clusters. All differentially expressed genes from each cluster were analyzed using EnrichR to determine the biological signature of the cluster.

In an independent analysis, Gene sets from KEGG, Biocarta, Reactome, and GO Biological Processes (MSigDB v7.2) were analyzed using GSVA on batch-corrected VST values, generating pathway activity scores (−1 to 1). Pathway scores were visualized over PaCMAP using ggplot2.

#### Copy Number Inference

Large-scale copy number alterations were inferred from bulk RNA-Seq using copykat. Manhattan plots were made using R package ggplot2.

#### Survival Analysis

Kaplan–Meier analyses were performed using samples with available recurrence data. Significance was assessed using the survival R package (v3.5.7). Multivariable Cox models were adjusted for relevant covariates as specified in Results.

#### Gene set variation analysis

Gene set variation analysis (GSVA) was used to quantify pathway and signature activity at the sample level. Curated gene sets representing PAM50 subtypes, Lehmann TNBC programs, CAF states, TAM states, immune programs, neuronal-like signaling programs, cell-cycle and DNA replication pathways, and therapy-associated response or resistance signatures were compiled from published studies and pathway databases, and are listed in Table S2. For each gene set, GSVA was applied to the log2(TPM+1) counts to generate an enrichment score for every sample.

#### Oncoscape Integration

Expression and clinical matrices were formatted for cBioPortal compatibility and uploaded into Oncoscape with predefined visualization states corresponding to manuscript figures.

#### Projection of external patient samples onto the reference landscape

External patient samples from GSE254461 were projected onto the breast cancer reference landscape to evaluate whether newly profiled tumors could be positioned within established transcriptional states. Expression data from the 19 external samples were processed to match the reference matrix by retaining shared protein-coding genes and applying the same normalization format used for landscape construction. Samples were then landed onto the reference embedding using two complementary approaches. First, we used the open-source PaCMAP transform function to project external samples into the existing PaCMAP coordinate space without refitting the reference landscape. Second, we applied a laboratory-developed nearest-neighbor projection approach, in which each external sample was matched to transcriptionally similar reference samples and assigned to the corresponding region of the landscape.

Concordance between the two projection strategies was assessed by comparing the landing position and cluster-region assignment for each external patient sample. PAM50 subtype scores were also calculated for each projected sample using the same subtype-scoring framework applied to the reference cohort. Projected sample positions were then compared with their calculated PAM50 subtype assignments to determine whether basal, HER2-enriched, Luminal A, and Luminal B tumors landed in the expected regions of the reference landscape.

## Supplemental Figures

**Supplementary Figure 1.**
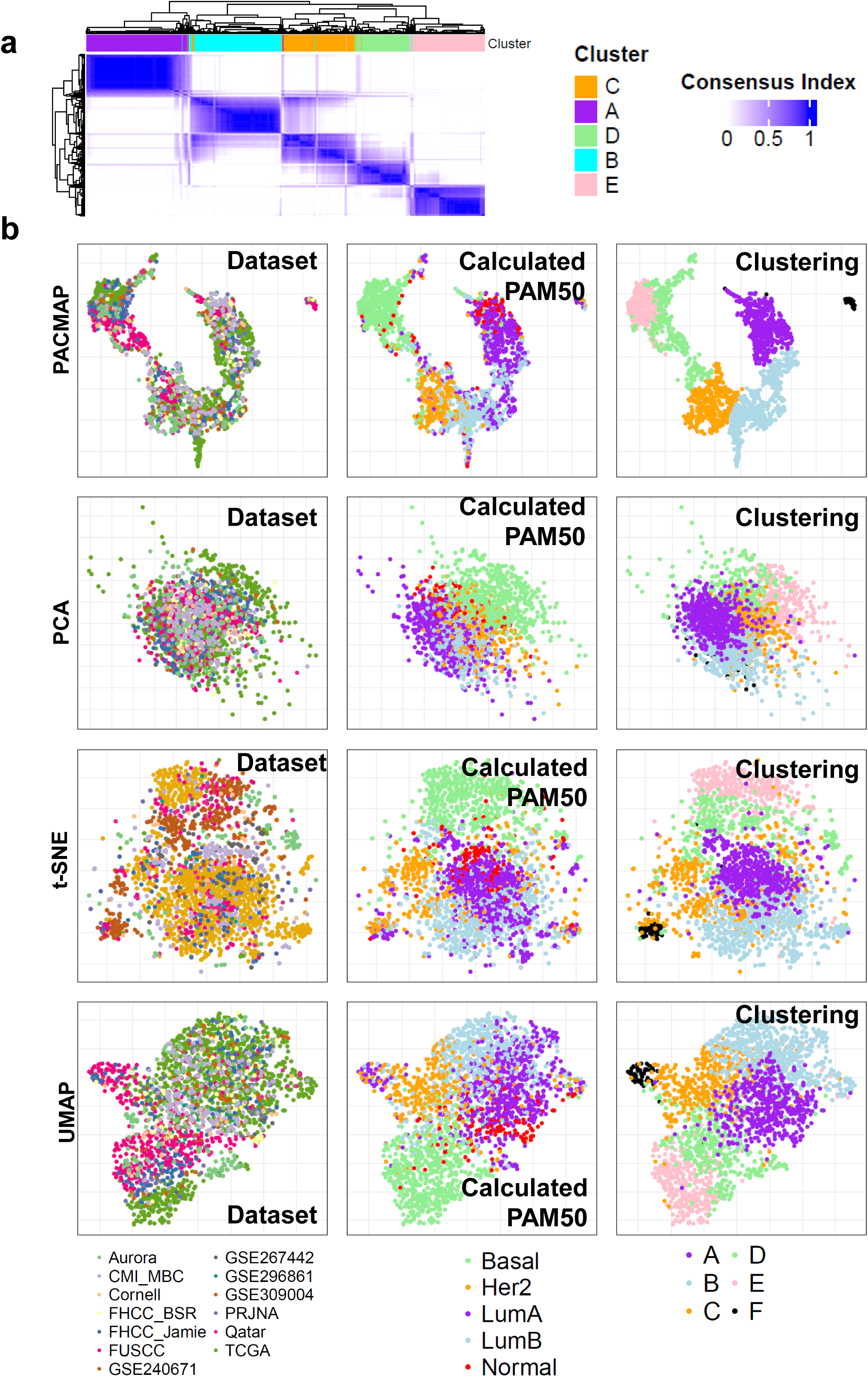
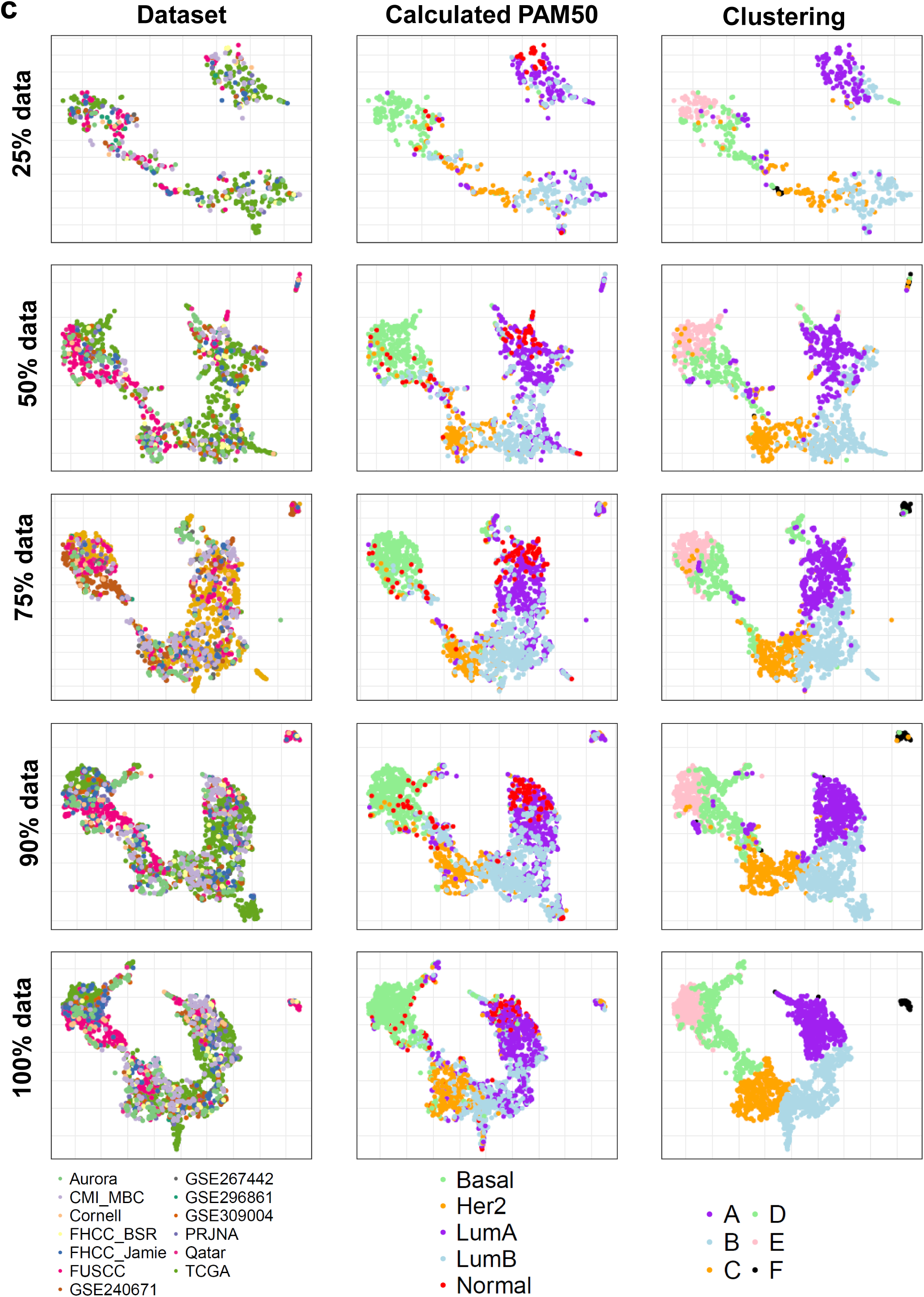
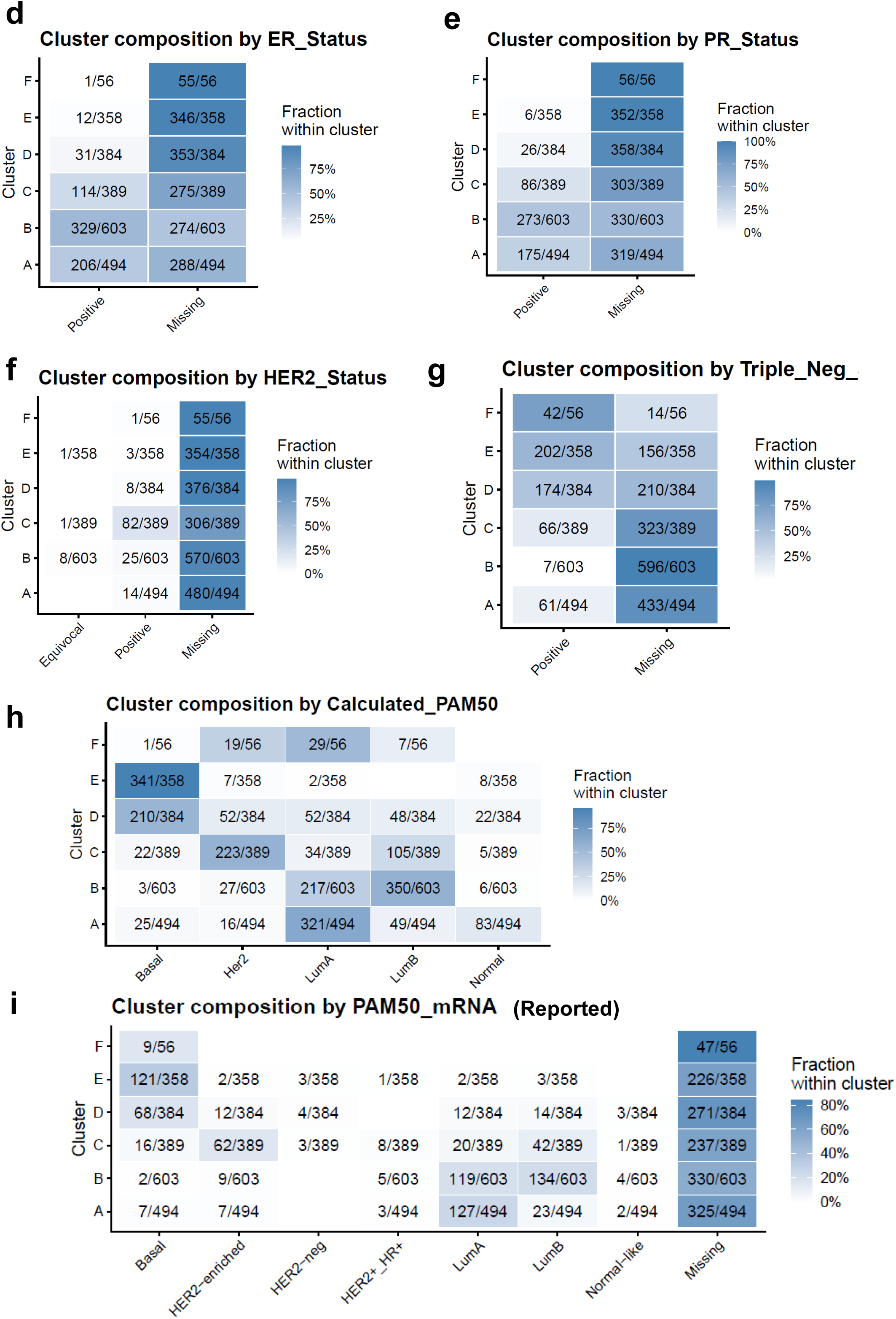
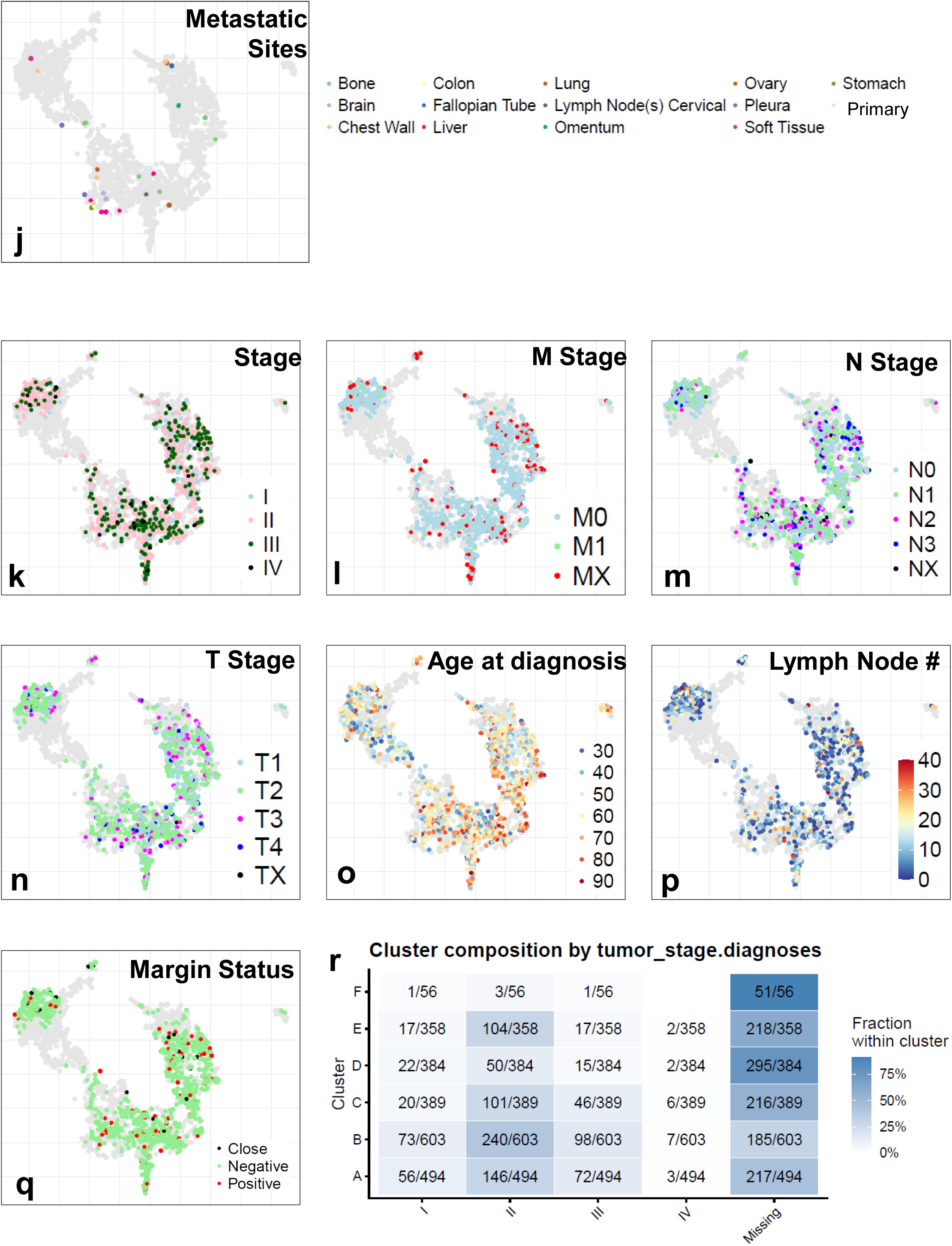
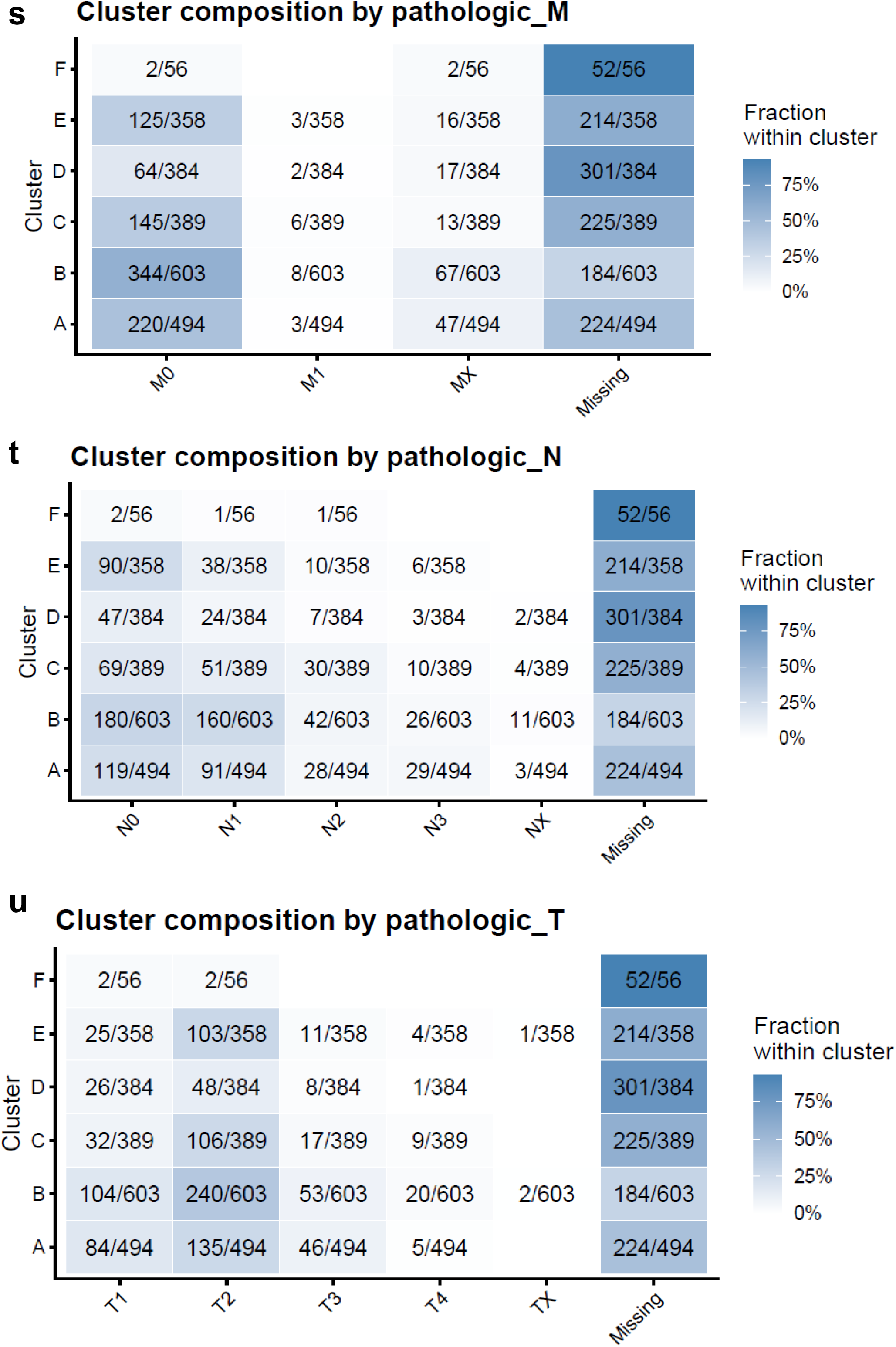
Landscape construction, visualization, robustness analyses and clinical annotation summaries. (A) Consensus clustering of the integrated breast cancer transcriptomic cohort using all protein-coding genes, identifying five stable transcriptional clusters. (B) Comparison of dimensionality-reduction methods used for visualization. PCA, t-SNE, UMAP, and PaCMAP embeddings are shown, each colored by dataset of origin, calculated PAM50 subtype, and consensus cluster assignment. Clustering was performed independently of dimensionality reduction; embeddings were used only for visualization. (C) Subsampling analysis assessing robustness of the breast cancer landscape. PaCMAP embeddings were regenerated after randomly sampling 25%, 50%, 75%, and 90% of the integrated cohort, with each embedding colored by dataset of origin, calculated PAM50 subtype, and consensus cluster assignment. Preservation of global structure across subsampled datasets supports the stability of the landscape. (D–I) Heatmap summaries showing the number of samples per cluster for ER status, PR status, HER2 status, triple-negative status, calculated PAM50 subtype, and reported PAM50 subtype. (J) PaCMAP embedding colored by metastatic site annotations for samples with available metastatic-site metadata. (K–Q) PaCMAP embeddings colored by additional clinical variables, including tumor stage, pathological M stage, pathological N stage, pathological T stage, age at diagnosis, lymph node status, and margin status. (R–U) Heatmap summaries showing the number of samples per cluster and margin status.

**Figure S2.**
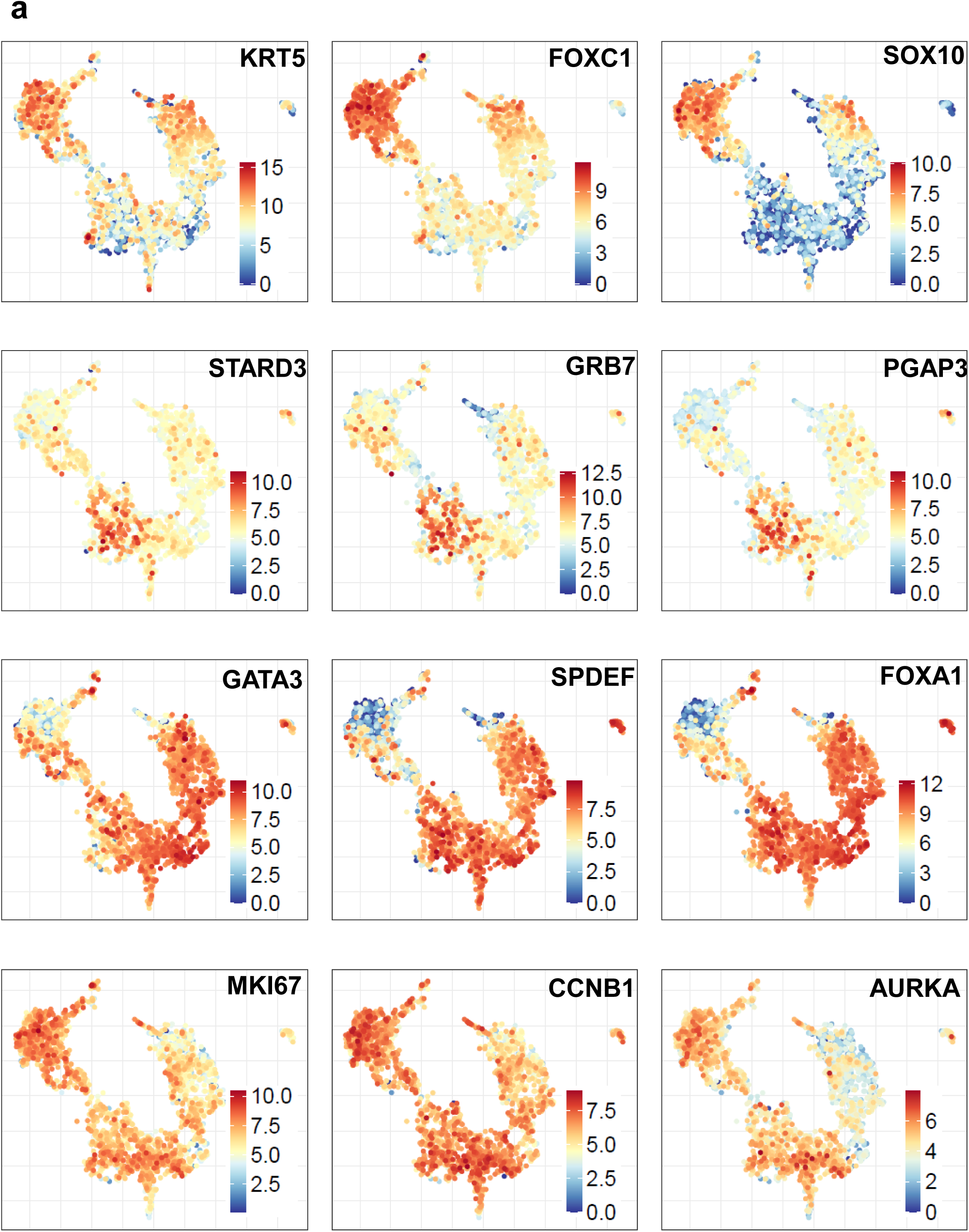
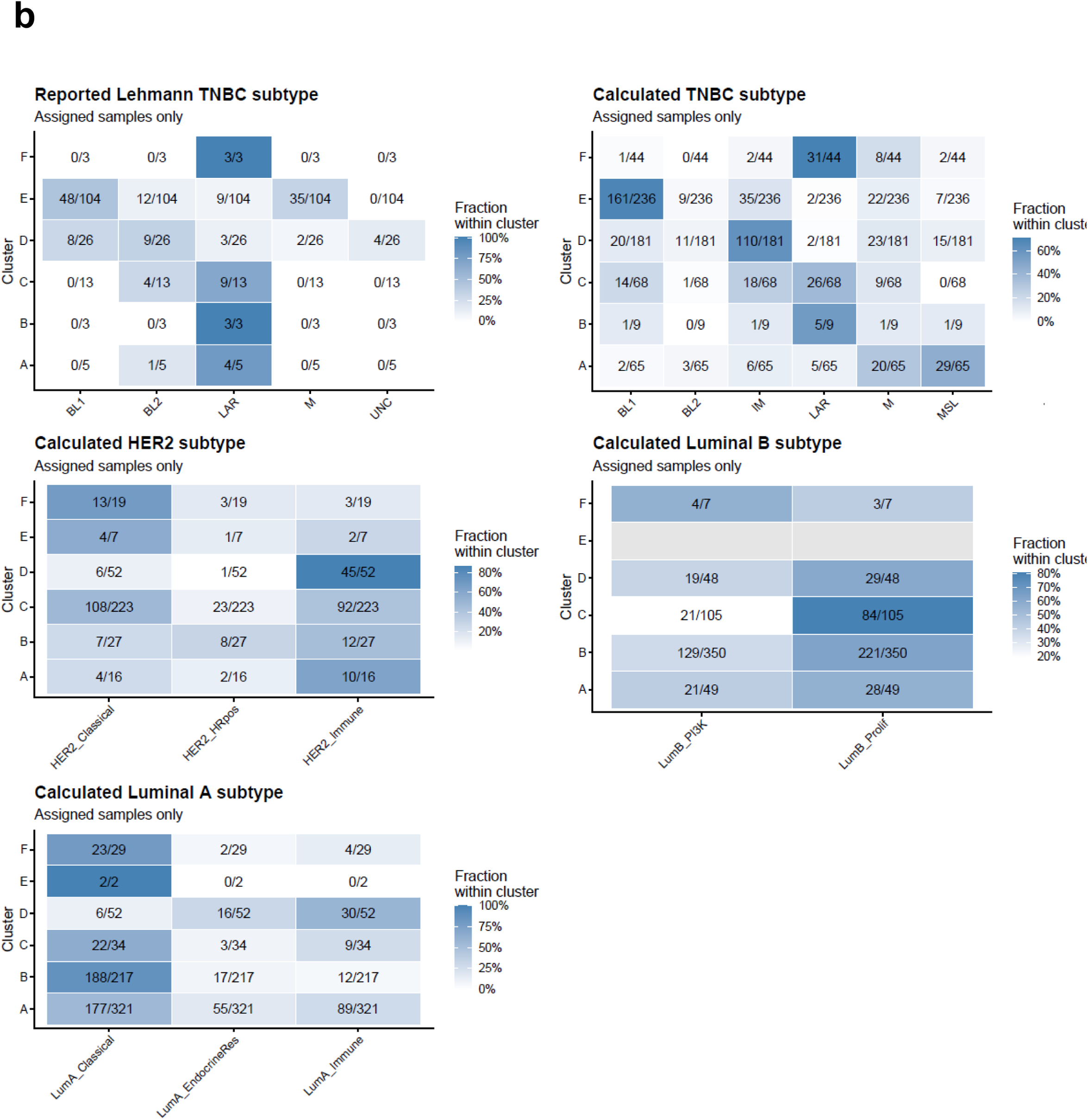
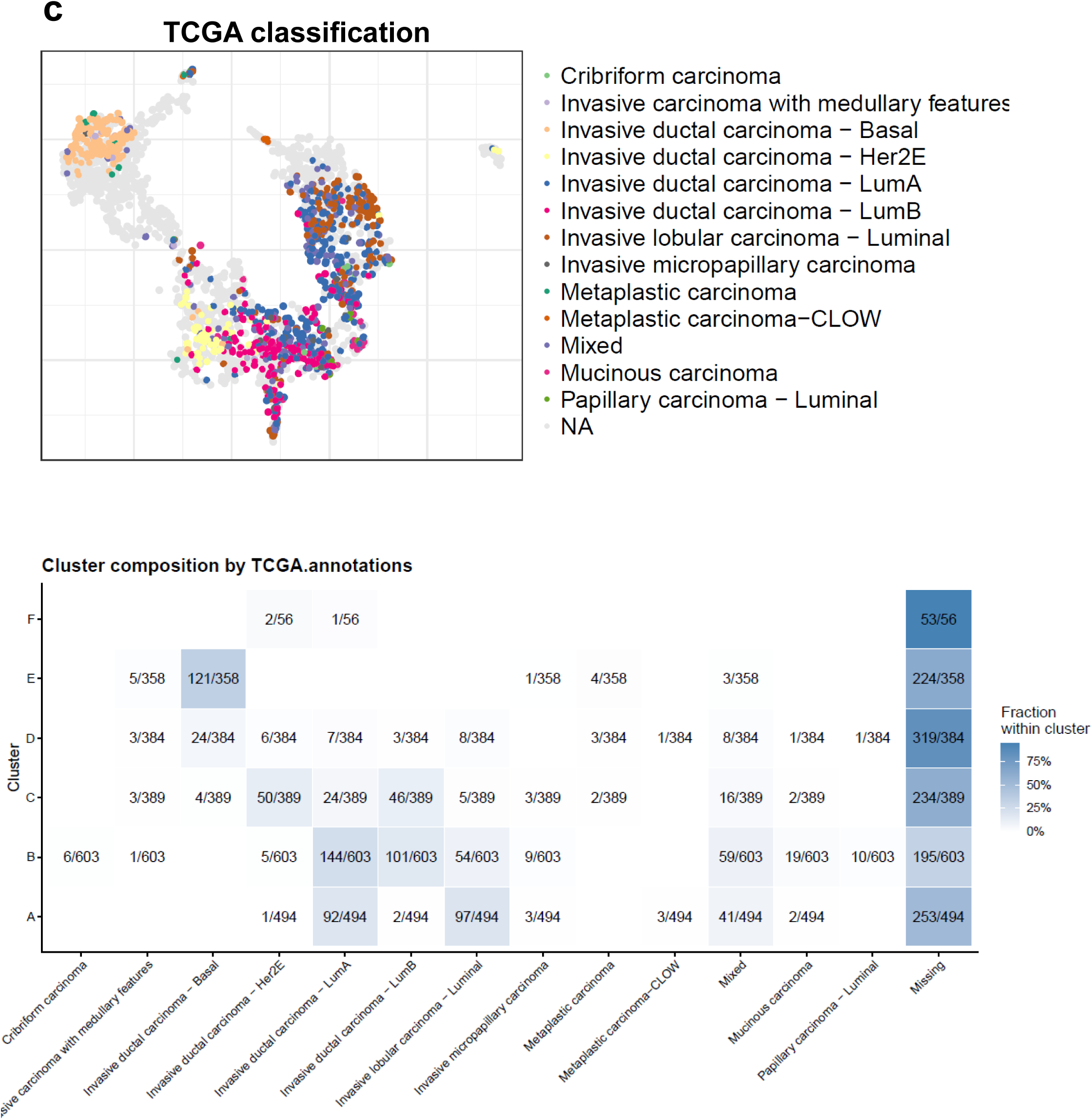
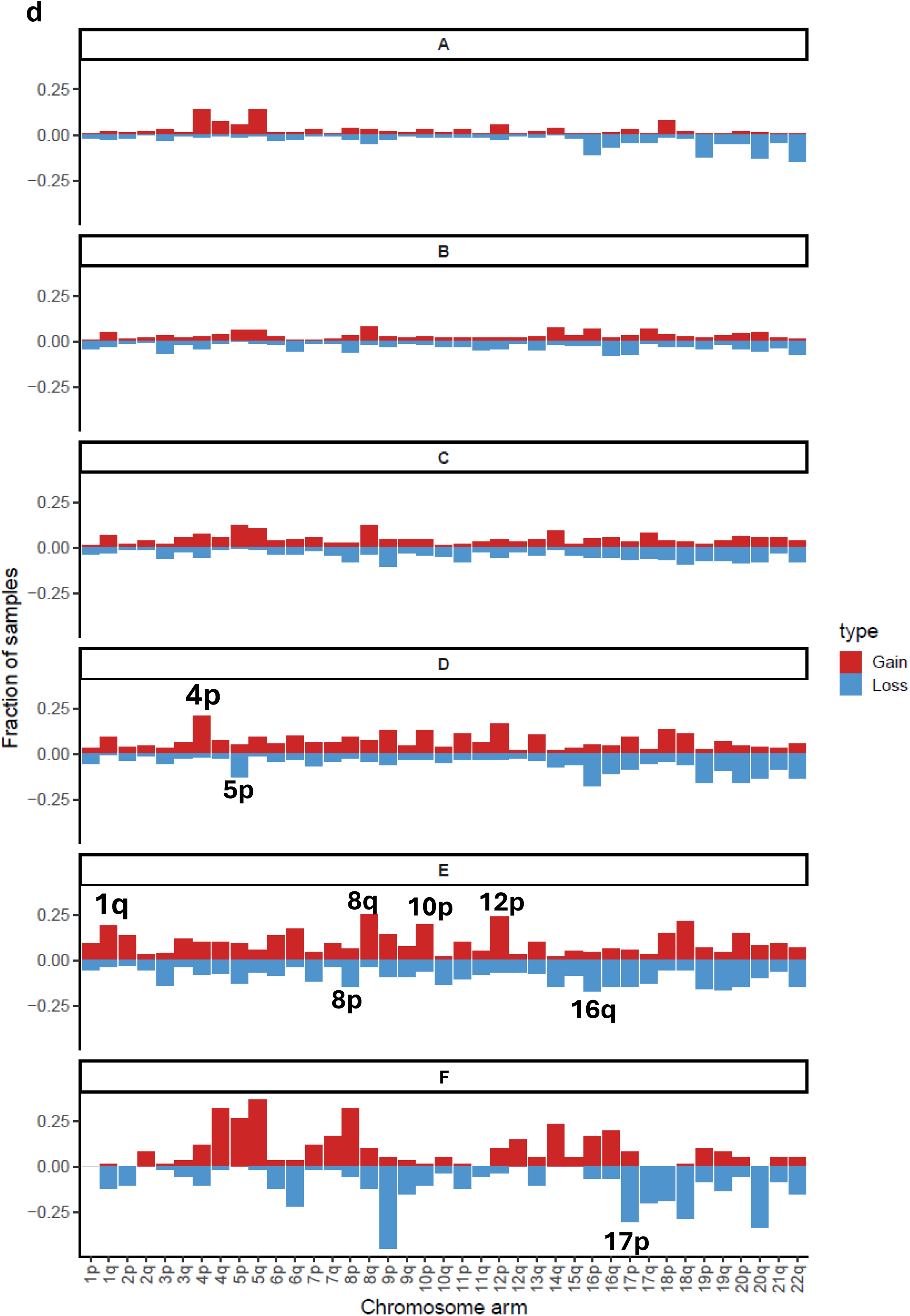
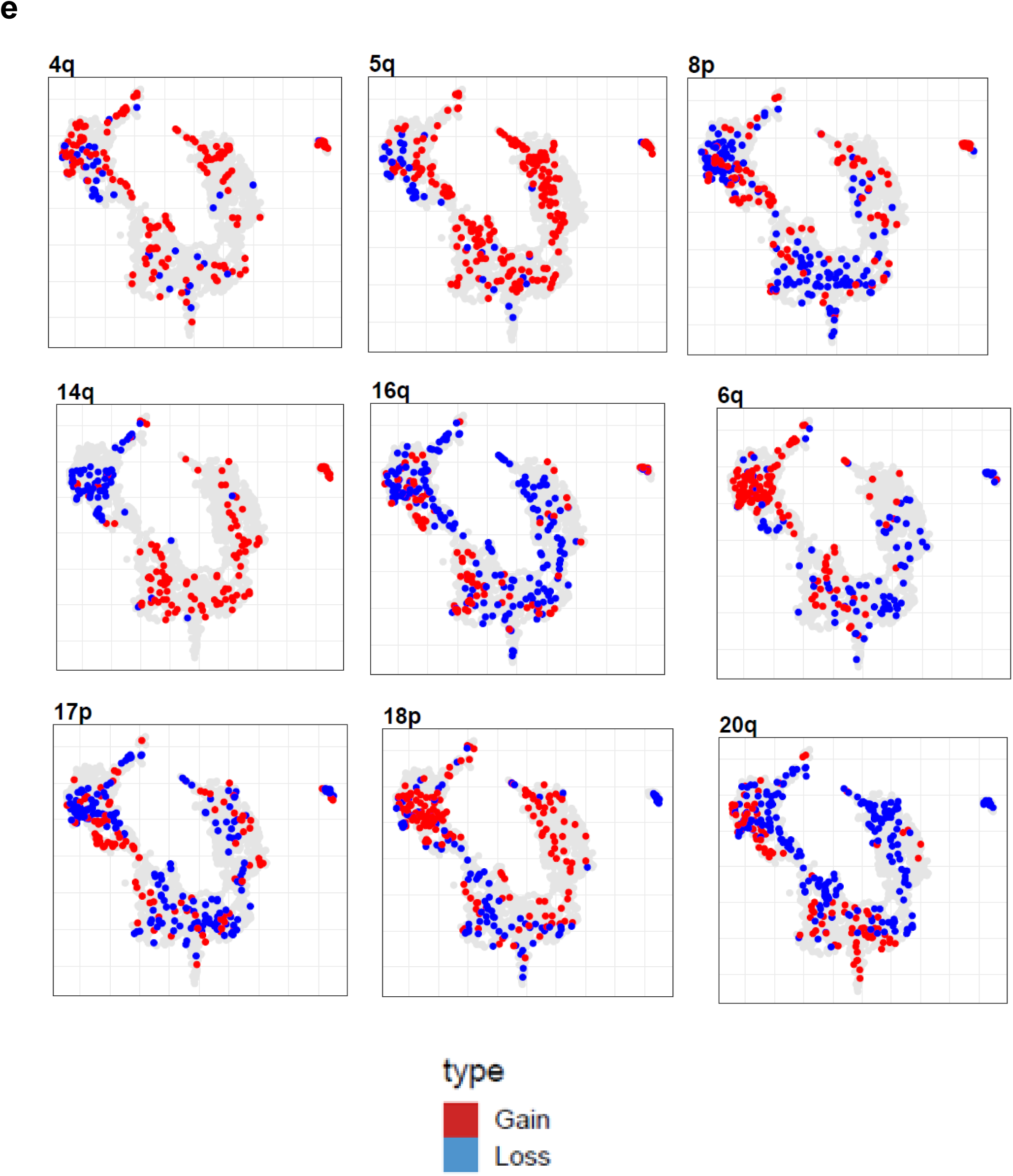
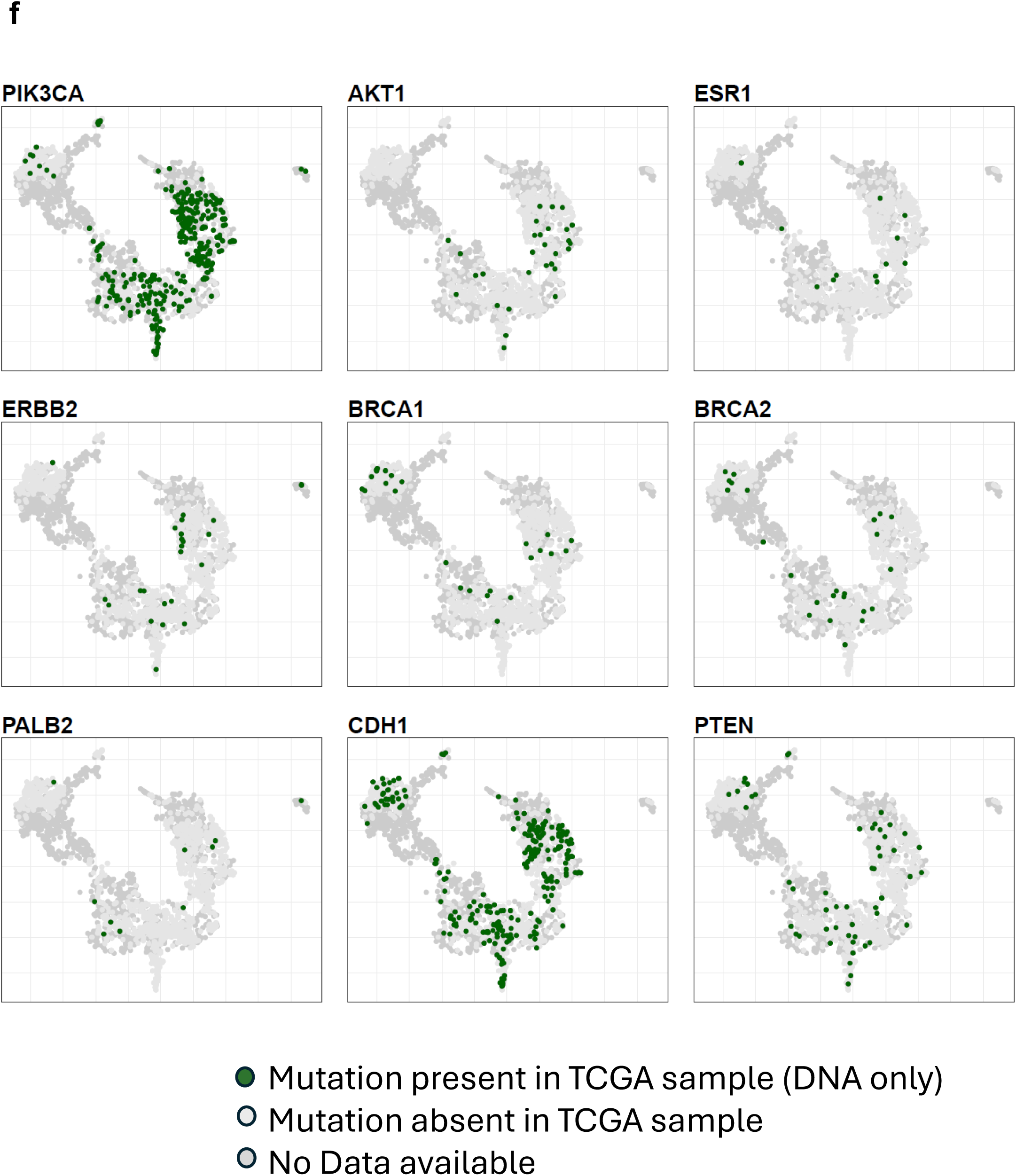
Gene expression markers define subtype-specific regions. (A) Expression of basal markers (KRT5, FOXC1, SOX10) enriched in the basal region. Expression of HER2-associated genes (STARD3, GRB7, PGAP3) localized to the HER2-enriched region. Expression of luminal markers (GATA3, SPDEF, FOXA1) enriched in Luminal A regions. Expression of proliferation-associated genes (MKI67, CCNB1, AURKA) enriched in Luminal B regions. (B) Heatmap summaries showing the number of samples per cluster for TNBC, HER2 and Luminal A/B subtypes. (C) Heatmap summaries showing the number of samples per cluster for TCGA biological subtypes. (D) Manhattan-style copy-number plots showing recurrent chromosomal arm-level gains and losses across landscape clusters. Each row represents one cluster or region, enabling comparison of cluster-specific copy-number alteration profiles. (E) PaCMAP landscape colored by selected chromosome arm-level copy-number alterations, showing the spatial localization of recurrent gains and losses across the transcriptional landscape. (F) PaCMAP landscape colored by DNA mutations for select genes

**Figure S3.**
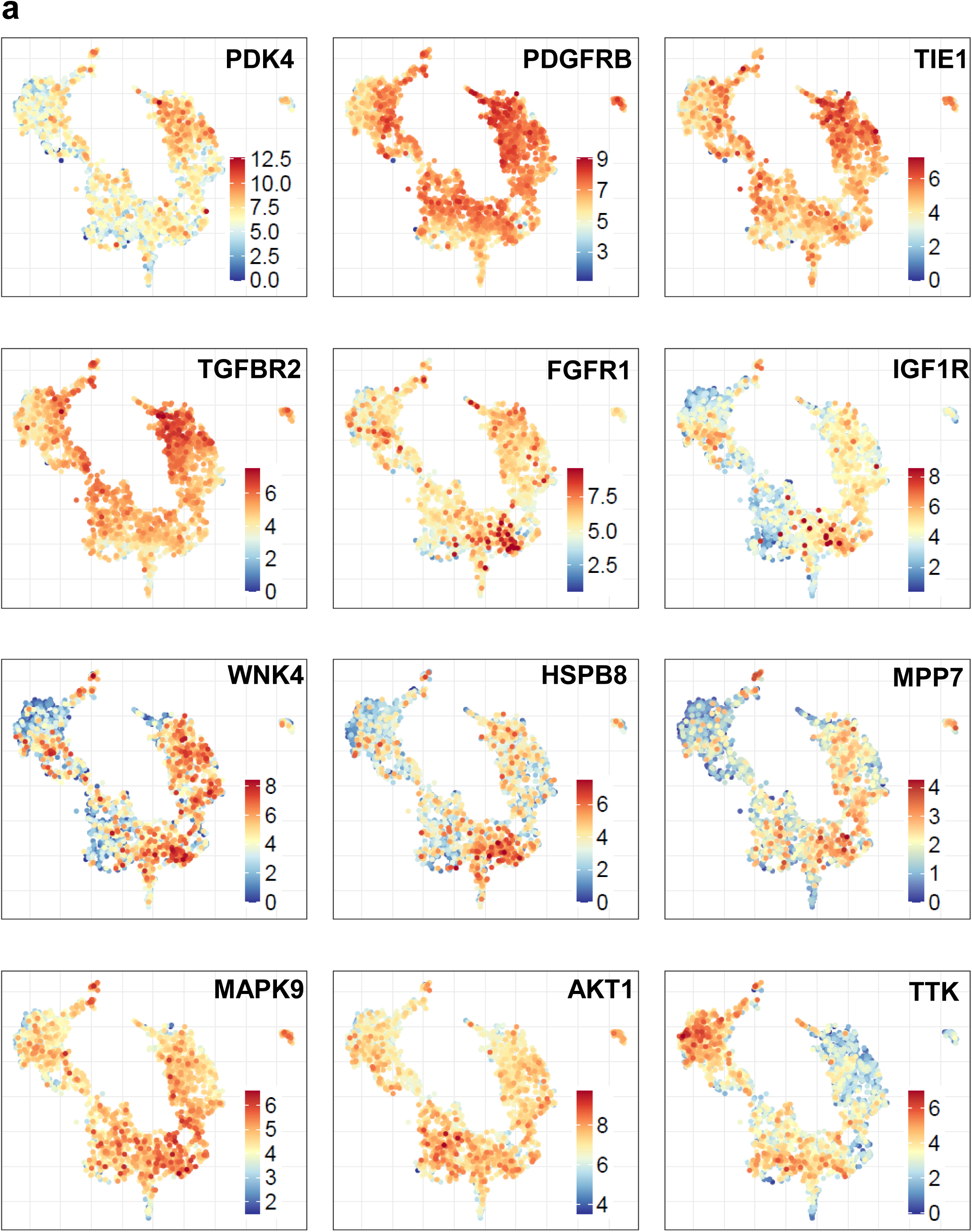
Gene and kinase expression programs across landscape clusters. (A) PaCMAP plots showing expression of representative kinase-associated genes enriched across different landscape clusters. These projections highlight cluster-specific kinase and signaling programs, including luminal-associated, HER2-associated, and basal-associated signaling states.

**Figure S4.**
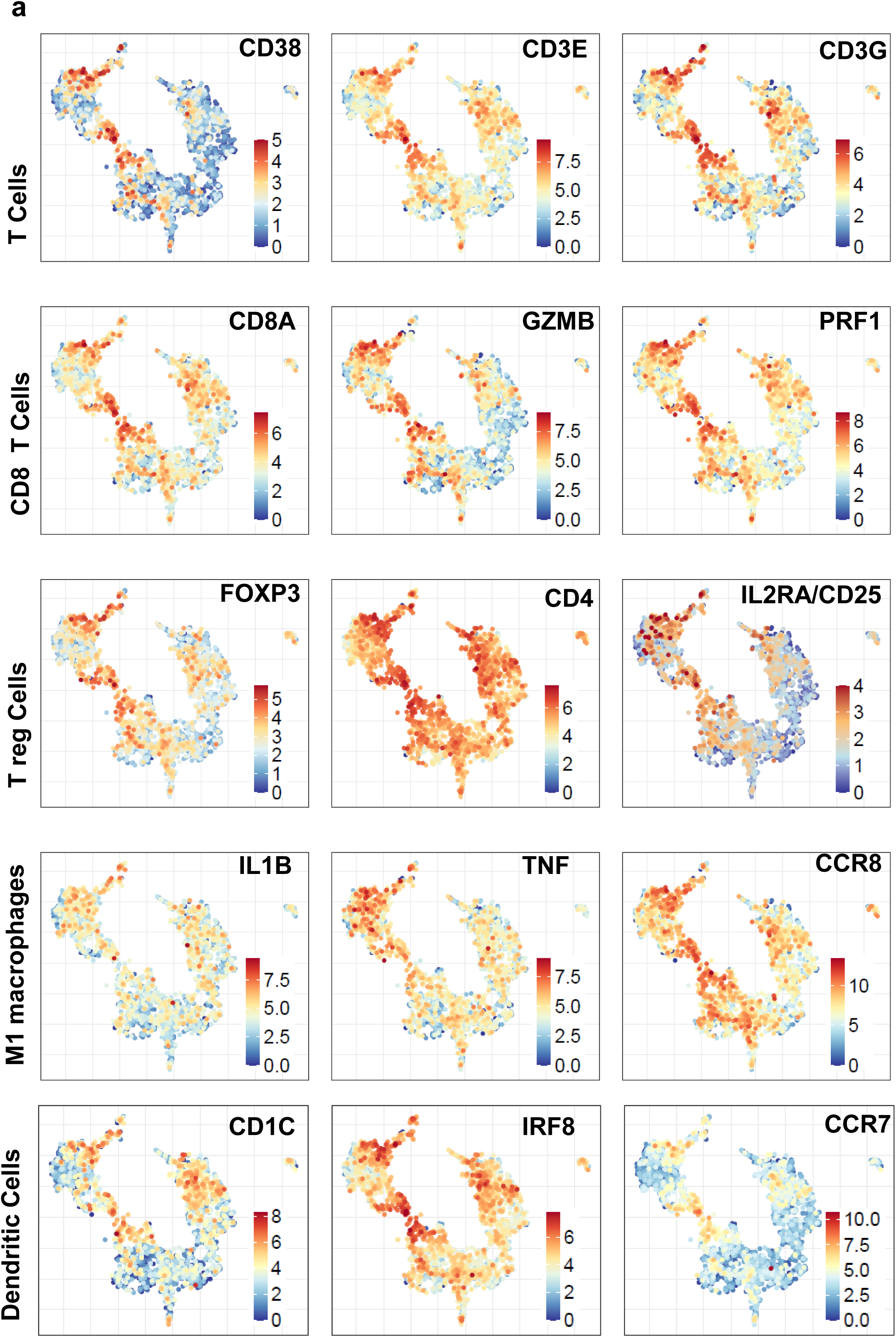
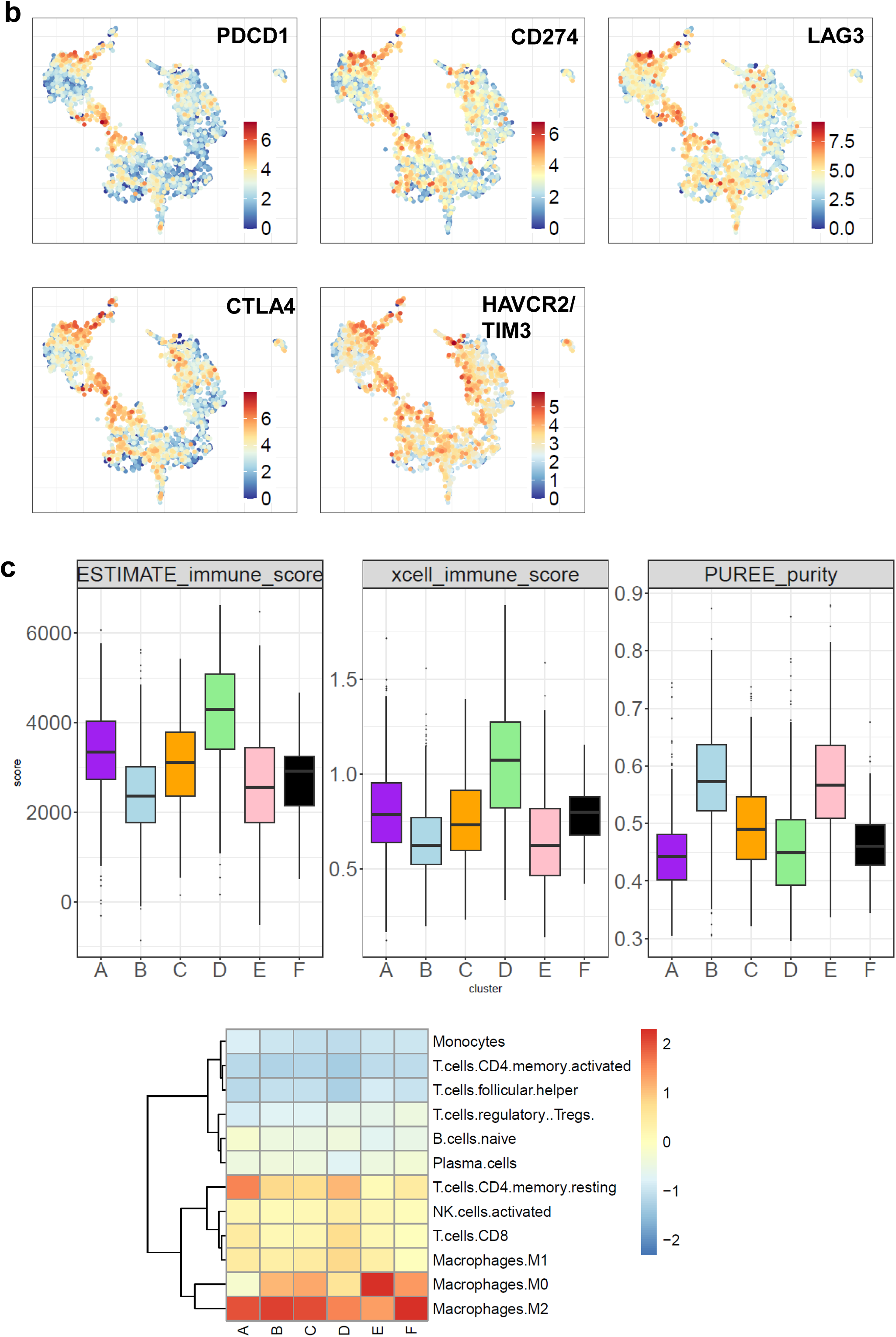
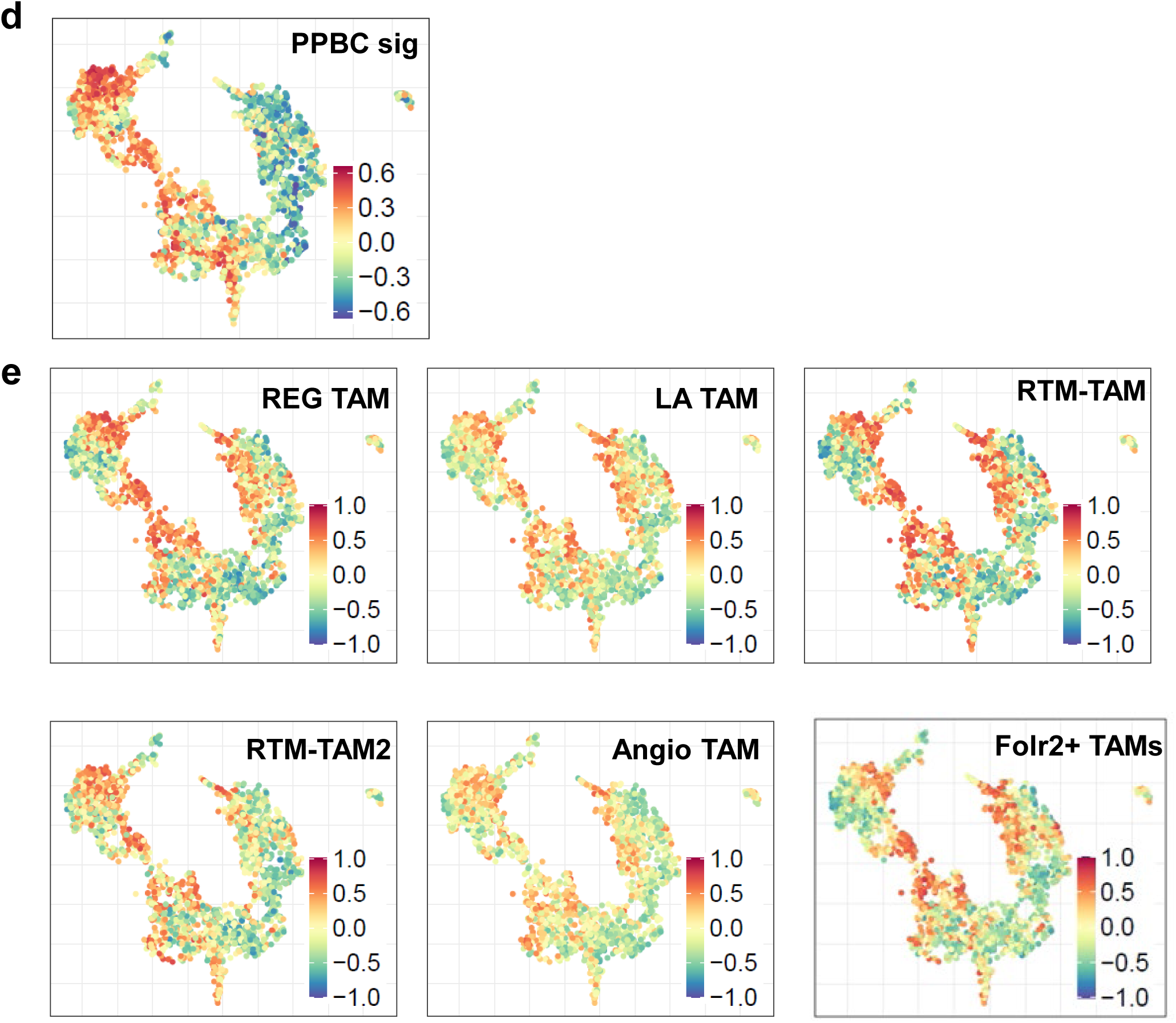
Immune and TAM-associated programs across the landscape. (A) PaCMAP plots showing immune cell–associated signatures, including T cells, CD8 T cells, regulatory T cells, M1 macrophages, and dendritic cells. These signatures are enriched predominantly in the immune-enriched basal-associated region. (B) PaCMAP plots showing expression of immune checkpoint genes PDCD1, CD274, LAG3, CTLA4, and HAVCR2/TIM3, supporting coordinated checkpoint pathway enrichment in immune-active regions of the landscape. (C) Boxplots showing immune score ( ESTIMATE and xCell) and tumor purity per cluster. Heatmap showing median cibersortX LM22 scores for each cluster. (D) PaCMAP projection of the postpartum breast cancer (PPBC) GSVA score, showing enrichment of this aggressive cell-cycle and immune-associated program across basal-associated regions. (E) PaCMAP projections of TAM-associated GSVA scores, including regulatory TAM, lipid-associated TAM, resident-like TAM, resident-like TAM-2, and angiogenic TAM programs. These signatures reveal spatial heterogeneity of macrophage states across the landscape.

**Figure S5.**
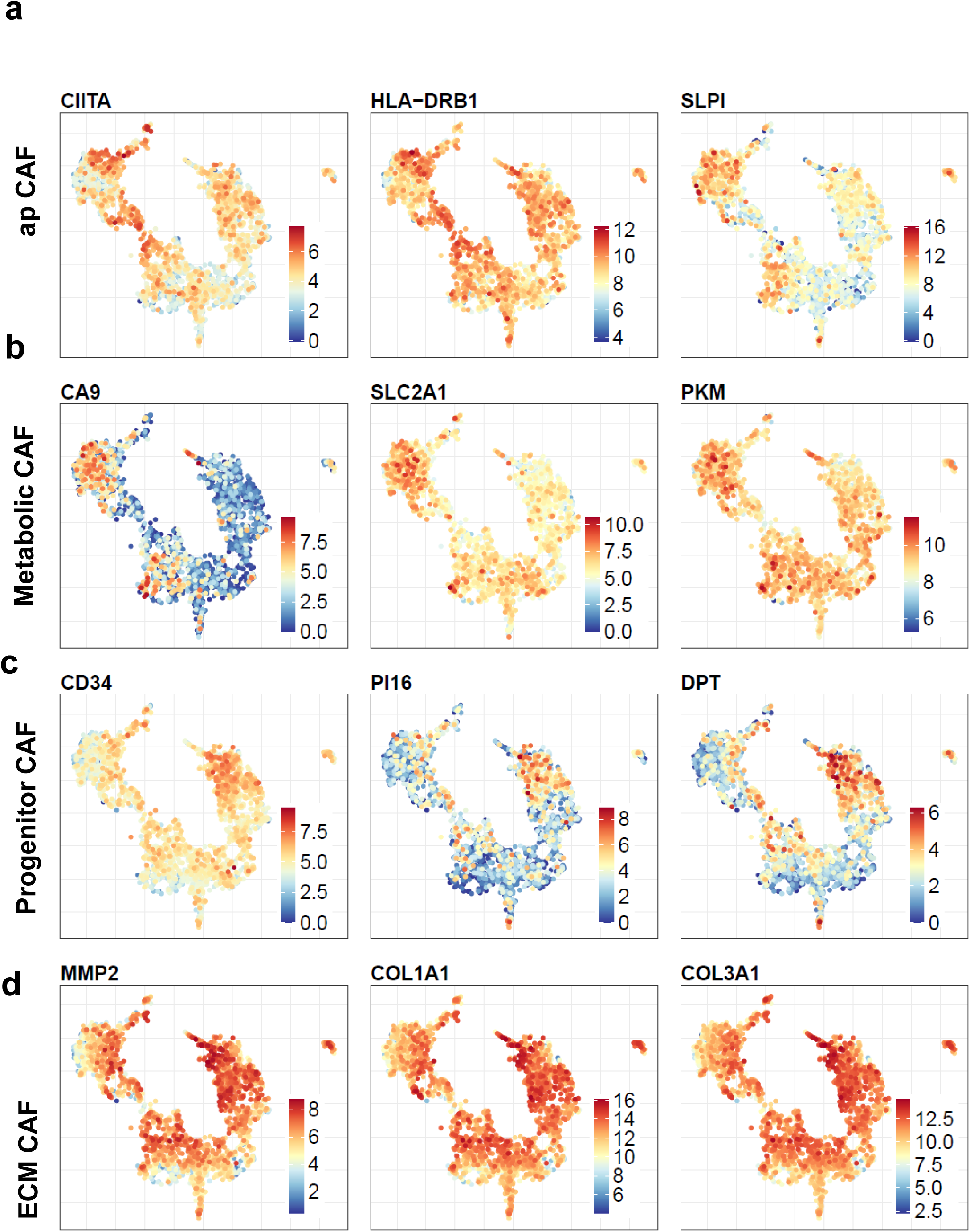
Marker gene expression supports CAF-state assignments. PaCMAP plots showing expression of representative marker genes for CAF programs used in GSVA-based CAF-state analysis. **(A)** apCAF-associated markers CIITA, HLA-DRB1, and CD274. **(B)** Metabolic CAF-associated markers CA9, SLC2A1, and PKM. **(C)** Progenitor CAF-associated markers CD24, PI16, and DPT. **(D)** ECM CAF-associated markers MMP2, COL1A1, and COL3A1.

**Figure S6.**
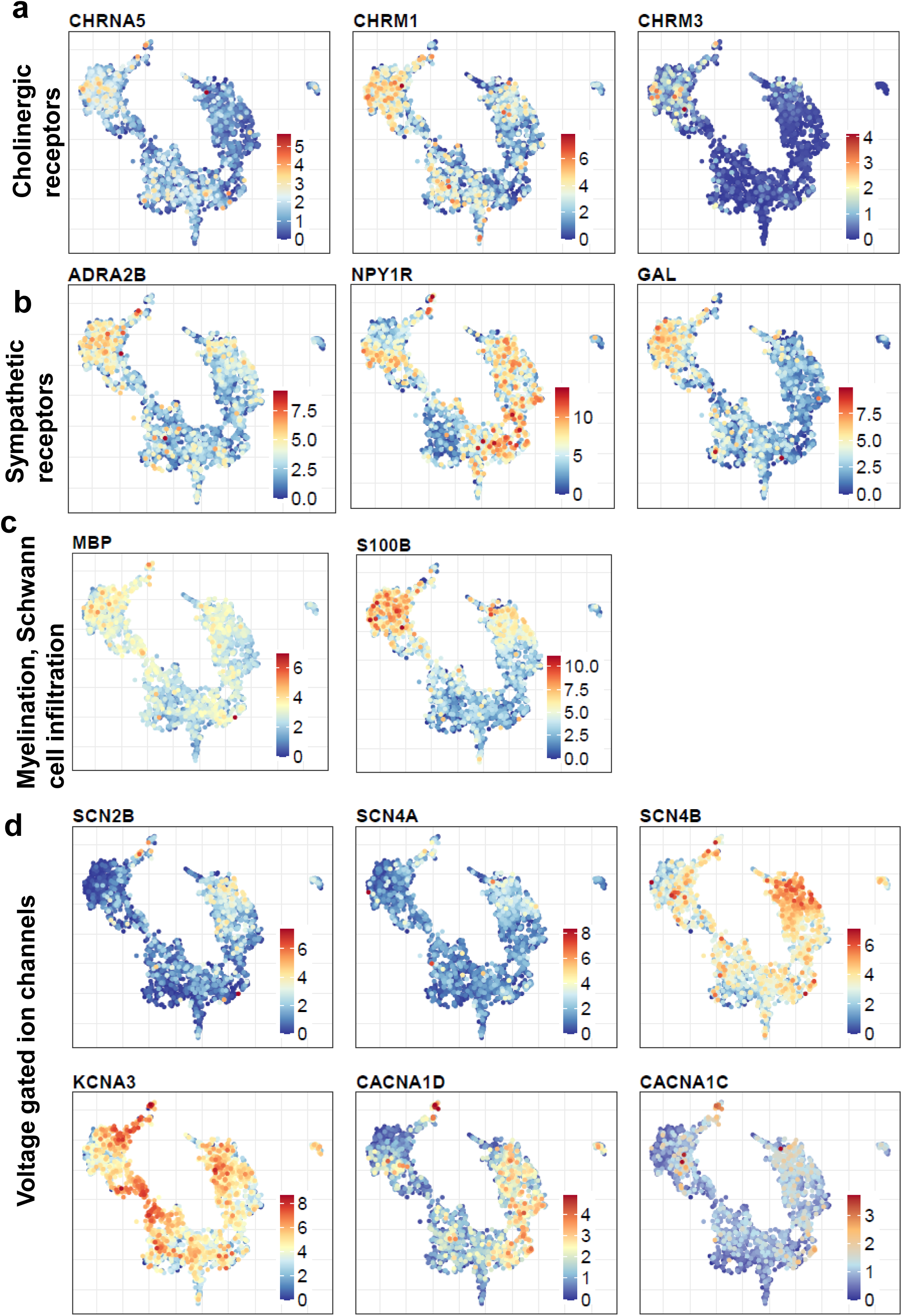
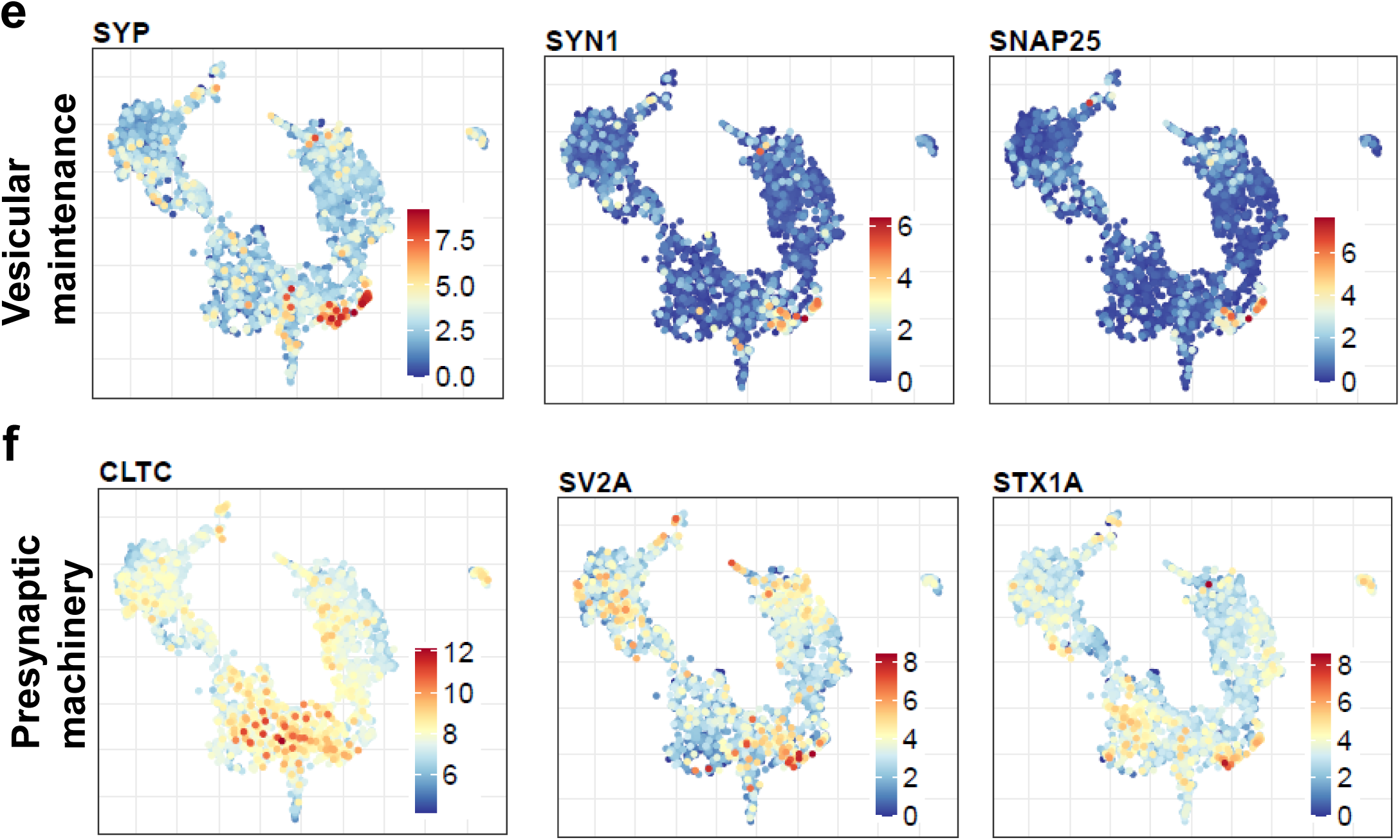
Neuronal-like signaling gene expression across the landscape. PaCMAP plots showing expression of neuronal-like and synaptic-associated genes across landscape clusters. (A) Cholinergic receptor genes CHRNA5, CHRM1, and CHRM3. (B) Sympathetic signaling-associated genes ADRA2B, NPY1R, and GAL. (C) Myelination-associated genes MBP and S100B. (D) Voltage-gated ion channel genes SCN2B, SCN4A, SCN4B, KCNA3, CACNA1D, and CACNA1C. (E) Vesicular maintenance genes SYP, SYN1, SNAP25 (F) Presynaptic machinery genes CLTC, SV2A, STX1A

**Figure S7.**
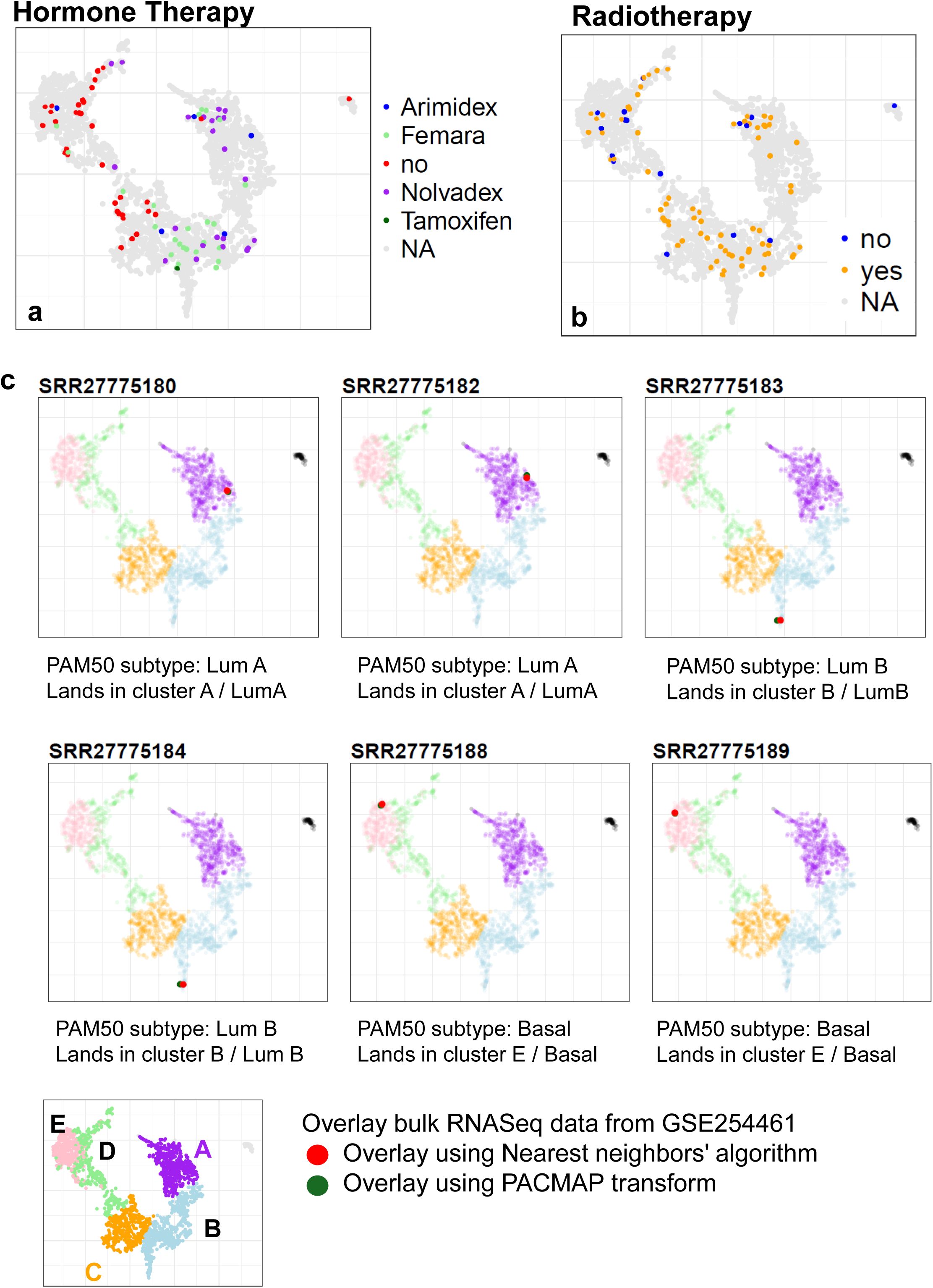
Treatment annotations across the breast cancer landscape. (A-B) PaCMAP projections showing TCGA treatment annotations for receipt of hormone therapy and radiotherapy. Treatment metadata were available for a subset of samples. (C) Comparison of external patient sample placement using two projection strategies: PaCMAP transform and a nearest-neighbor landing approach. Each of the 19 GSE254461 patient samples was projected onto the breast cancer reference landscape by both methods, with matching sample identities shown using the same color. Concordant landing positions between the two approaches support the robustness of patient projection into the reference landscape.

## Supplementary Table Legends

Table S1. Number of samples contributed by each dataset included in the integrated breast cancer transcriptomic landscape.

Table S2. Gene lists used for GSVA score for PAM50 subtype, TAM, CAF and drug response.

Table S3. Percentage for PAM 50 subtype calculation for each cluster

Table S4. Published subtype concordance and composition per cluster

Table S5. Chromosome arm–level copy number gains and losses stratified by clusters

Table S6. Differential gene expression analysis results across clusters.

Table S7. Enriched Pathways across clusters. Table S8. Kinases up-regulated in each subtype

Table S9. Synaptic Markers up-regulated in each subtype

Table S10. PaCMAP coordinates for landed patients on breast reference landscape.

## Acknowledgements

We thank members of the Holland lab at Fred Hutch Cancer Center for valuable discussions and collaborators for sharing their data and metadata. This research was supported by funding from the Fred Hutch Cancer Center (E.C.H), 1R35 CA253119-01A1 (E.C.H).

## Author Contributions

Conceptualization, S.A., and E.C.H.; Methodology, S.A. and E.C.H.; Formal Analysis, R.S. and S.A.; Software, M.J., G.G; Investigation, S.A. and E.C.H.; Resources, E.C.H.; Data Curation, S.A and R.S., Writing – Original Draft, S.A and E.C.H; Writing – Review & Editing, S.A., E.C.H S.H., H.P, Y.L.; Visualization, S.A., R.S., M.J., G.G; Supervision E.C.H., Funding Acquisition, E.C.H.

## Data and materials availability

All analysis including statistics and visualization were done in R version 4.3. Plots were generated using R basic graphics and ggplot2. Raw sequencing data was downloaded from GEO as shown in Table S1. All custom code used in this study is available at https://github.com/sonali-bioc/BreastLandscapes.

## Ethics

This study was conducted using publicly available and fully de-identified transcriptomic datasets. For new data collected from human subjects, institutional review board (IRB) approval and informed consent were required. All original studies from which data were obtained had received appropriate ethical approvals and consent for data sharing.

